# Structural dynamics and neural representation of wing deformation

**DOI:** 10.1101/2024.06.02.596338

**Authors:** Alexandra M. Yarger, Masateru Maeda, Igor Siwanowicz, Haruhiro Kajiyama, Simon M. Walker, Richard J. Bomphrey, Huai-Ti Lin

## Abstract

Locomotor control is facilitated by mechanosensory inputs that report how the body interacts with a physical medium. Effective representation of compliant wing deformations is particularly challenging due to the many degrees of freedom. Structural configurations can constrain the stimulus space, and strategic placement of sensors can simplify computation. Here, we measured and modelled wing displacement fields and characterized spatiotemporal encoding of the wing mechanosensors. Our data show how dragonfly wing architecture prescribes deformation modes consistent across models and measurements. We found that the wing’s state under normal flapping conditions is detected by the spike timing of few sensors, with additional sensors recruited under perturbation. The functional integration of wing biomechanics and sensor placement enables a straightforward solution for information transfer.

**Significance Statement:** Many systems in nature precisely control highly deformable structures, yet monitoring structural deformations has posed a significant challenge for biologists and engineers. By measuring and modelling the intricate structure of compliant dragonfly wings, we demonstrate an elegant solution for mechanosensory representation. We show that the functional integration of natural wing biomechanics and sensor placement provides a straightforward solution for information transfer. Wing morphology passively constrains the range of natural deformations, producing strain patterns that align with sensor locations, allowing them to monitor the wing using a simple timing-dependent encoding strategy. The way insects monitor aerodynamic and inertial forces via wing deformations during flight could apply to many sensory systems in nature and inspire artificial neural networks for controlling diverse dynamic systems.

## Introduction

To produce effective outputs during fast behaviours, where time constraints limit the amount of information that can be detected, nervous systems must have efficient and robust mechanisms in place for processing sensory inputs. An extreme example of this is the wing sensory systems of flying animals. Wings experience high loads at high speeds, which are complex but crucial for propulsion, stability and control. The unsteady aerodynamics of gliding and flapping flight interact with the wing’s structural properties, resulting in passive wing deformations. Dramatic wing deformation is characteristic of animal flight and has been tied to inertial loads, lift generation, steering, and stall prevention(1–3). But with so many degrees of freedom and dynamic states, how do flying animals extract sufficient information from aeroelastic wings in real-time? Insect wings are often used as a model to study this problem because there is no actuation beyond the wingbase, so deformation can be attributed to passive effects. Insect wings also have a large range of mechanosensors, with dragonfly wings being among the most sensor-dense (4, 5). Wing mechanoreceptors can also convey information from multiple sources, highlighting the potential complexity of wing sensory systems(6).

In the context of behaviour, it isn’t always necessary to represent every possible input condition constantly. In general, the precise tuning of insect sensors results from anatomy and neural encoding mechanisms that have evolved to extract only relevant information from the environment(7–13). Sensor tuning is modulated at multiple stages of the nervous system to allow dynamic and context-specific sensorimotor control(14–16). The stimulus space can be reduced *prior* to transduction by ‘morphological computation’, where information is filtered by structural mechanics, sensor location, or both. Sensor specificity is already well-documented for mechanosensation at the level of individual sensor morphology(17); for example, shape determines directional selectivity of strain-sensing campaniform sensilla (CS)(4, 18–21). Structural and location-based spatiotemporal filters for relevant mechanical inputs including cuticular strain, fluid flow, and sound are known across taxa(17–20, 22–25).

Similar mechanosensory encoding mechanisms and sensor distributions have been observed across insects(4, 5, 21, 26), but what are the features that determine optimal sensor locations? Thin rectangular plate wing models have previously identified sparse sensor placements for optimal representation of body rotations(27–29), but three-dimensional geometry is likely to have dramatic effects on deformations and hence sensor placement for optimal detection of the loading state during flight(2). To investigate this, we took a multipronged approach, measuring and modelling local and global deformation displacements and strains across the surface of the wings of tethered and freely flying dragonflies. We describe the effective stimulus space (the natural range of deformations that wings encounter) using dimensional reduction techniques and then characterize how only certain features are encoded (and are therefore available to the flight controller). Using information theory, we identify the most information-rich areas on the wings in relation to the actual sensor distribution (30, 31). With this synthesis (summarized in Movie S1), we demonstrate an elegant mechanism of morphological computation for mechanosensory representation in a biological system. The general principle, based on insects’ ability to monitor instantaneous aeroelastic loading conditions via deformations during flight, is likely to be applicable across sensory systems in nature and could form the basis of guidelines for the design of sensor arrays for artificial neural networks used to control diverse dynamical systems.

## Results

### Free-flight control

Steering control has been extensively studied in insects(32–36). However, wing elevation, sweep, and pitch are typically the only kinematic variables considered, despite documentation of considerable variation in deformations, consistent with a role in flight control(2, 3, 37). To demonstrate that wing deformations are significant and correlated to flight maneuvers, we used a nine-camera, high-speed, free-flight arena to measure wing kinematics throughout the stroke cycles of freely flying dragonflies (*Sympetrum striolatum*; Fig. 1A). We found that even relatively straight flight trajectories result in differences in the patterns of deformation between wing strokes; for example, bend amplitude (displacement from resting state = 0) varied between the two forewings, though the phase relationship between them remained stable (Fig. 1B-D).

**Fig 1.**
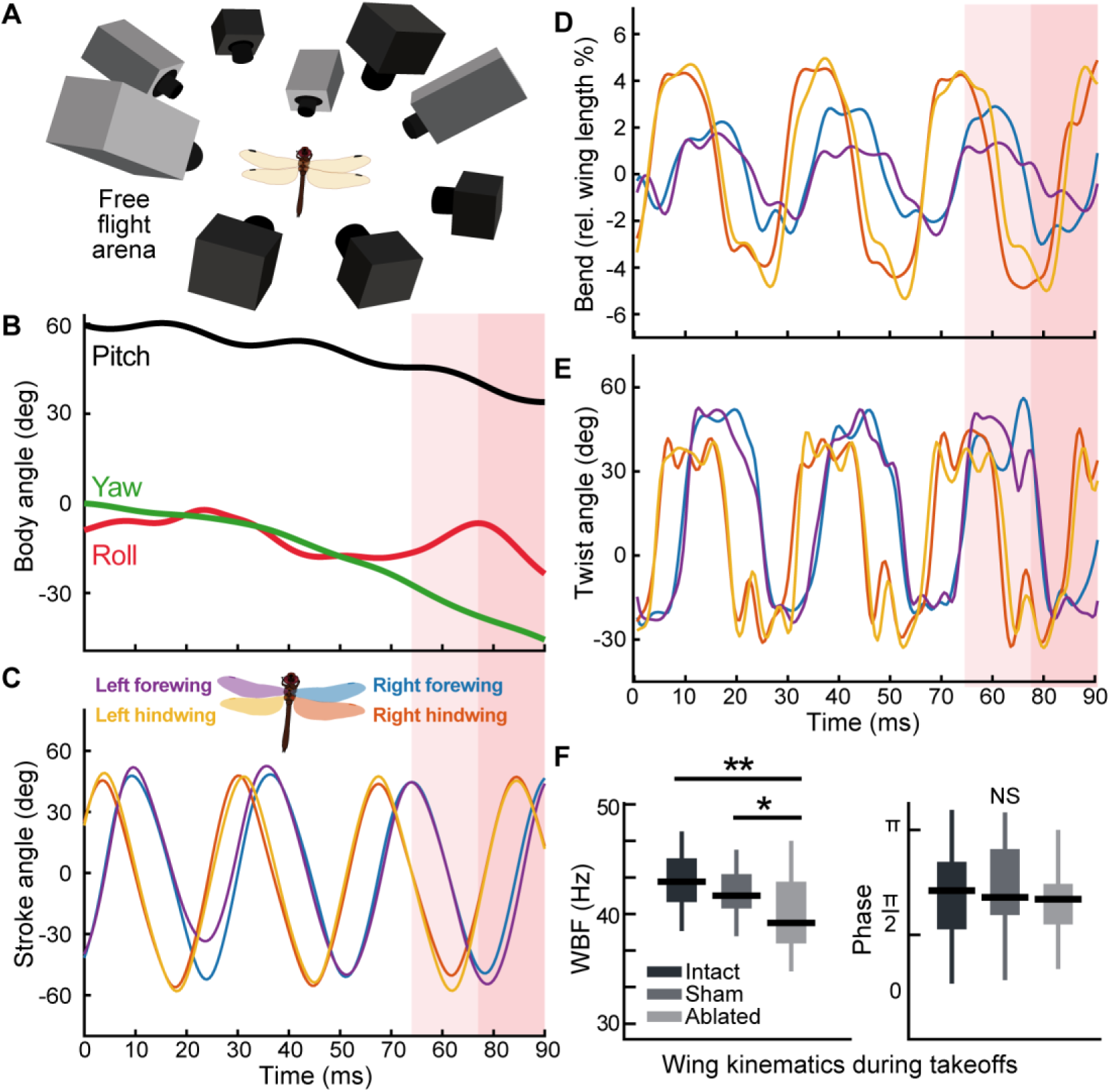
Free-flight kinematics. A) Nine synchronised high-speed cameras captured free-flight wing deformations at 2000 frames per second (fps). B) Body orientation over time of freely flying dragonfly (see Movie S2). Orientation is relatively straight for initial two wingbeats. Roll manoeuvre highlighted in red occurs during the third stroke cycle. C) Wing stroke angle (calculated relative to mean stroke plane) showing comparative wing timing across three wingbeats. D) Wing bend amplitude change across wing strokes. E) Forewing twist pattern changes during roll manoeuvre. F) Ablated wingbeat frequency (WBF) was significantly lower than intact (P=0.00002) and sham (P=0.0024) conditions during takeoffs filmed with one camera at 1000 fps. There was no significant difference between intact and sham WBF, and no significant phase difference between all conditions (see table S1).

During a rolling turn (highlighted), there were changes to the deformation patterns compared with straight flight; and clear asymmetries in the left and right forewing twist and bending patterns (Fig. 1E). Given the mechanosensory array at their disposal, these results suggest that strain patterns would be encoded differently by each wing’s sensory system.

To determine whether wing sensors are important for flight control, in a separate experiment, we ablated the wing sensors by lesioning the anterior wing nerves. These nerves contain the wing mechanosensory afferents and project to wing motor neurons (Fig. S9B). We measured how sensor ablation affected the animals’ ability to perform take-offs - a key flight maneuver that has been described in detail and can be broken down into stereotypic components(38). Here we focused on the first four full wingbeats, during which time the fore- and hindwings move in synchronous phase. Nerve-ablated take-off attempts resulted in lower elevation gains than nerve-intact and sham-ablation take-offs (Fig. S1A). Nerve-ablated dragonflies also frequently abandoned take-off attempts once airborne and would glide or fall to the ground (Movie S3).

Within individual conditions (either intact, sham, or nerve-ablated), there were no significant differences between each consecutive wingbeat (Fig. S1), so all four wingbeats were pooled for comparisons between conditions. There were no differences in forewing-hindwing phase between all conditions and there were no differences in wingbeat frequency (WBF) between intact and sham. However, nerve-ablated animals did have significantly lower WBFs compared to both intact and sham animals (Fig. 1F). This is consistent with a previous study where it was shown that the central nervous system contains all the neuronal machinery required to produce the flapping motion but sensory feedback (including information from the wings) is required to maintain appropriate flapping frequency(39). Having confirmed that wing sensors are necessary to maintain frequencies that generate adequate lift during take-off, we moved on to identifying inputs that the wing sensors are monitoring.

### Wing deformation patterns

To understand how dragonflies monitor wing structural deformations, we first characterized the natural range of wing deformation patterns using three parallel experimental approaches. Each approach was designed to measure displacement of the wing relative to the wingbase reference frame, ensuring that deformation within the wing was captured independently from displacements due to wing hinge articulation (i.e. flapping). First, we drove a wing into aeroelastic flutter(40) using an unsteady wind stimulus. The fluttering wing, immobilized at the hinge, was filmed using a high-speed camera. A digital image correlation (DIC; see Methods) analysis was then used to reconstruct the wing’s displacement field (Fig. 2B). Second, we re-processed the wing deformation from the nine-camera free-flight data in the same format as the flutter displacement field maps (Movie S4). Finally, we produced a high-fidelity wing model incorporating fine details of vein geometry for finite element modal analysis (FEA) and fluid-structure interaction (FSI) gliding simulations (Fig. 2C). FEA also yielded high-resolution strain fields, which we later use to assess the efficacy of the CS strain sensor distribution (Fig. 4G).

**Fig 2.**
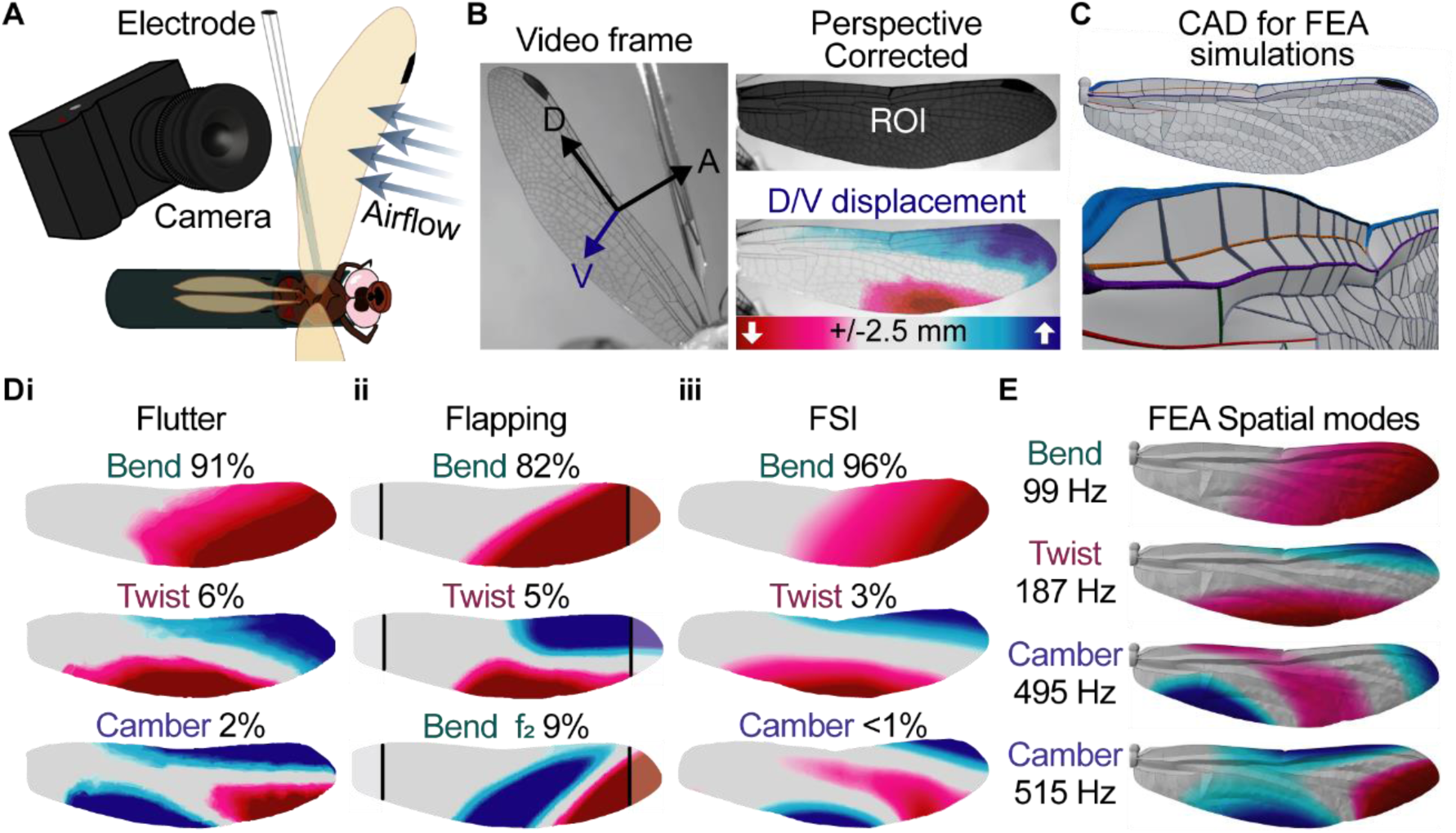
Wing deformation parameters. A) Wings were mounted in fluctuating airflows. Surface deformations were filmed with a high-speed video camera at 1000 fps and the neural activity of the wing mechanosensory afferents were recorded simultaneously. B) Wings were filmed at an angle to maximize visualization of dorso-ventral (V) displacements (distal-proximal (D) and anterior-posterior (A) displacements were negligible; see Fig. 3S2A). After perspective correction, digital image correlation (DIC) was used to measure displacement. C) A 3D external morphological model of a dragonfly forewing reconstructed by hybridizing the live wing vein pattern (determined by direct linear transformation of stereoscopic views) with vein cross-sectional geometries (isolated from μCT scans; top). Bottom shows rotated close-up view. D) PCA results. i. Deformation modes during flutter, representing 99% of overall variance (N=5 animals, 14,000-49,000 frames/animal). ii. Deformation modes during free flapping flight (base/tip interpolated from measured displacements) representing 96% of overall variance (F_2_=2^nd^ harmonic). iii. FSI (15°, 2.6 m/s) deformation modes representing 99% of overall variance E) FEA deformation modes ranked by oscillation frequency (see Movie S5). FEA camber modes have similar patterns & frequencies (dominated by deflections in both directions along the trailing edge) and are likely represented together within camber mode of the fixed wing preparation flutter experiments and FSI simulations.

To characterize the natural wing deformations, we sought to represent the overall wing motion with the fewest necessary variables by dimensional reduction. A principal component analysis (PCA) of the fluttering wing data revealed that 99% of the wing’s displacement variance can be described by the first three components, interpreted as bending, twist, and camber structural modes (“Flutter”, Fig. 2Di). The same modes were also present in our flapping flight measurements (“Flapping”, Fig. 2Dii) except for the mode representing camber, which is not currently measurable as the method only reconstructs the leading and trailing edges. However, camber has previously been documented in free-flight in dragonflies(41) and is a known essential feature of insect wing deformation(2, 35, 42). Nevertheless, the bend and twist modes represented 96% of the displacement variance. PCA of FSI simulation (Movie S1) revealed the same modes (“FSI”, Fig. 2Diii), and they also emerged as natural modes in FEA simulations (Fig. 2E) demonstrating that our high-fidelity wing geometry model captures the physics of deformation (Fig. S2). Together, they show that a linear decomposition of wing displacements reveals a handful of modes inherent to the wing structure that are excited during gliding and flapping flight. Could the wing sensory system leverage the structural properties to encode the wing deformation using a handful of modes? We investigated this possibility by characterizing the mechanosensor encoding mechanisms.

### Temporal selectivity

Does the sensory array detect the spatial modes excited during flight? To test this, we recorded neural activity simultaneously with airflow stimulation (Fig. 2A). The wing deformed periodically in response to the airflow and the frequency of deformation contains information about the system’s mechanical behaviour(43–46). We anticipated that deformations could be phasically encoded and looked for phase information in afferent signals passing into the thoracic ganglia through the wing’s anterior nerve. A template-matching custom spike sorting application(47) was used to separate the responses of sensory neurons into individual ‘units’ within each tethered recording; typically, recordings contained 1-3 distinguishable units. To identify the primary deformation features driving each unit, we first performed PCA on the high-dimensional wing displacement data to extract the most significant modes for each unit. Spike-triggered averages (STA) were then computed and visualized along these PCA-derived modes (Fig. 3A), providing an intuitive representation of the temporal response while preserving the underlying multidimensional feature structure. Covariance analysis also confirmed that PCA accurately captures the dominant response features (Fig. S4D). The recorded sensors are sparsely activated and cannot be recorded for long periods due to their proximity to the mechanical stimulation. As a result, applying nonlinear models such as maximum noise entropy or maximally informative dimensions led to overfitting and unintuitive results. In general, STA combined with simple feature identification methods (e.g. PCA) are sufficient for describing signals in peripheral neurons, especially mechanosensors(48), and serve the purpose of this study. Due to the high dimensionality of our data, we performed feature extraction prior to STA. However, covariance analysis of the spike-triggered raw displacements was also performed on the trial shown in Figure 2 and two features were identified: bend and twist (Fig. S4D).

**Figure 3.**
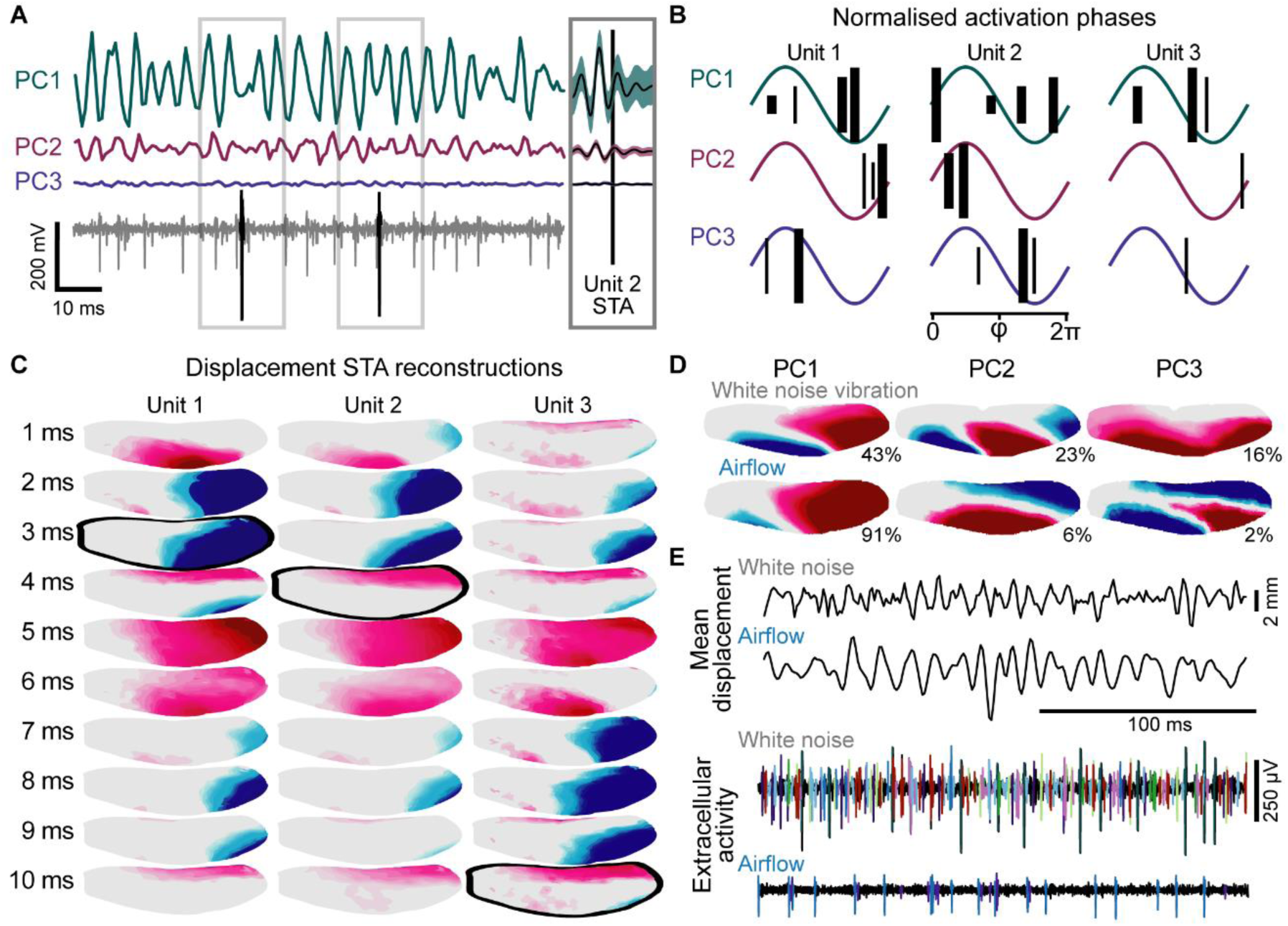
Phase encoding. A) Principal component (PC) displacement scores and the associated afferent spike response to airflow-induced wing deformations were used to calculate spike-triggered averages (STA) for each unit (see also Fig. S4A). Vertical black bars represent spike times for unit 2. B) Activation timing of phase-locked units for each PC (See Fig. S4B). Vertical bars show spike time relative to PC oscillation. Thick and thin bars represent strong (vector strength>0.6) and weak (0.6>vector strength>0.5) phase-locking, respectively, and bar height corresponds to displacement amplitude (1-4 mm). Phase timing shifts with displacement amplitude. C) Heatmap reconstructions of PC1-3 STA for airflow induced deformations with 4 mm mean absolute displacement of the entire wing. Each image represents a single frame (1 ms). Images highlighted in black show the spike time. D) White noise vibrations result in novel displacement modes (colormap as in Fig. 2). E) White noise induced displacements (above) result in greatly increased neural activity compared to airflow induced displacements (below). Colours indicate different units.

Across trials, the 1-2 largest units (signal to noise ratio>4) dominated the overall activity of the recording and were consistently phase-locked to the bending and twisting modes (and, less strongly, to camber) at the wing’s natural frequency (Fig. 3B). It should be noted that these modes are always present simultaneously and were only separated here to simplify analysis and allow 2D visualization of the STA. When all three features are recombined, we still find that these units responded to the same patterns of deformation at different phases (Fig. 3C); the phase of activation for these units was observed to shift depending on the displacement magnitude (Fig. 3B). These responses enable the dragonflies to encode derived conditions: for example, asymmetries between contralateral wings, or bending mode magnitude at different points within the stroke cycle. To avoid misrepresenting responses to cyclical stimulation using STA, units that were not phase-locked were excluded in the analysis (Fig. 3C). Phase-locked units are defined by vector strength (Fig. S4)(49, 50). Importantly, in our tethered preparation the wing base is fixed, so inertial rotations of the wing relative to the body are absent. This indicates that the phase-locked neural responses we observe are driven by aeroelastic deformations arising from aerodynamic and structural forces, rather than by inertial or gyroscopic effects.

An important feature of biological control systems is that they should be robust to novel stimuli. To understand how wing sensors respond to deformations outside the observed spatial modes and frequencies, we presented unnatural perturbations using a lever arm attached at a single point, the nodus, vibrating with uniform white noise (0-500Hz). Vibration-induced displacements were confirmed to be outside of the range observed in our tethered and free-flight experiments by comparing spatial modes (Fig. 3E) and frequencies (Fig. S6) to those previously identified (Fig. 2E). These differences were also reflected in the neural signal; white noise vibration resulted in increased spike frequency, and the recruitment of additional units at lower displacement amplitudes (Fig. 3F). In airflow stimulus recordings, all large units were phase-locked (vector strength>0.5) with good signal to noise ratio (>2). These units were also activated in response to lever-arm stimuli, but reliable unit identification was not possible due to additional overlapping units that emerged. Though these new units could not be characterized here, our results suggest that the large number of strain sensors may be critical for monitoring unnatural perturbations.

### Spatial selectivity: sensor localization

Extracellular recordings do not allow direct identification of the neurons. However, due to the causal effect, the mechanosensor activity should maximally correlate to the wing area where the strain sensors are located. To identify candidate sensors, we calculated coherence between the activity of each unit and the displacement or strain within small spatial bins (akin to pixels) across the wing surface (Fig. 4A). Coherence is a measurement of the correlation between two signals across frequencies and represents how well one signal predicts another. High coherence represents more accurate predictions, in this case how well a unit predicts the wing’s deformation or vice versa. The coherence and peak coherence frequency of a single unit and the displacement of the wing across spatial bins is shown in Fig. 4A-C.

**Figure 4.**
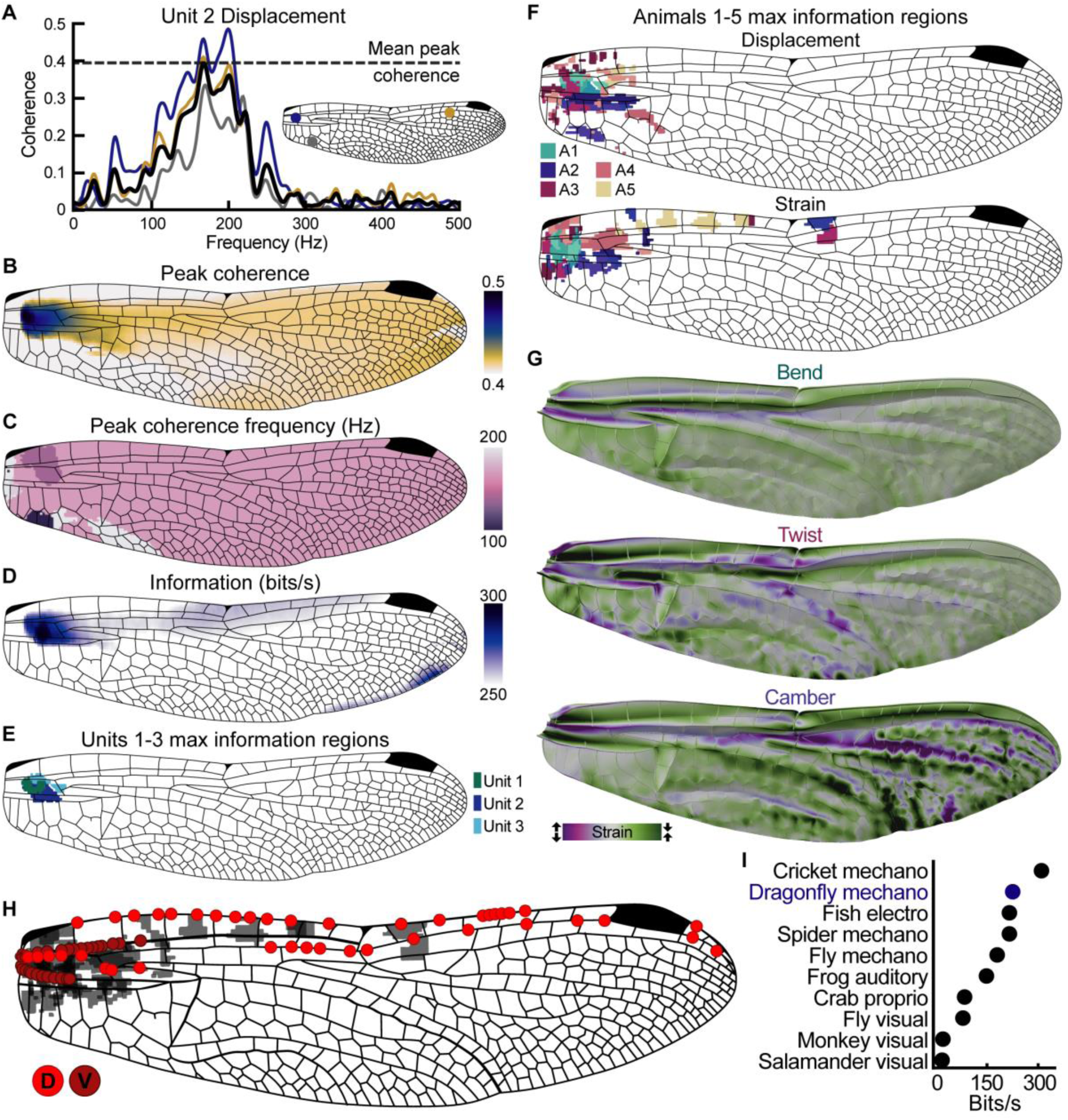
Sensor localisation. A) Coherence between the activity of a single sensory unit and the displacement of the wing for three example spatial bins and the mean peak coherence for all bins across the wing surface (black) between 0-500Hz. Unit 2 peak coherence for the mean of all bins is 171Hz. B) Peak coherence values of all bins. Darker regions show higher coherence. C) Frequencies at which the peak coherences shown in panel B occurred. For this unit, most peaks occur at 171Hz. D) Information rate of unit 2 estimated from coherence between 0-500Hz of all bins using the lower bound method. E) Wing regions with the highest normalized displacement information rates for each of the recorded units previously shown in Fig. 3. The darkest region in D corresponds with the dark blue region shown here (unit 2). F) Wing regions with the highest normalized displacement (top) and strain (bottom) information rates across animals (N=5 animals, n=3,3,2,2,1 cells respectively). Colours correspond to individuals and shade variations show different units recorded from the same individual. Data in A-E are from animal 1 (A1). G) FEA spanwise strain for key deformation modes (green-compressive, purple-tensile) showing regions of high strain. H) Locations of CS on major veins. Dorsal and ventral denoted by marker colour. Shaded regions show high strain information rates from F. Sensors located in these regions represent likely candidates for the recorded units. I) Lower bound estimates of information rate for dragonfly CS (mean of all cells’ maximum rate across the wing surface) compared with other spiking sensory systems(48, 84–92).

Coherence was high across a range of frequencies, so to capture the stimulus-response correlation for all frequencies, we converted coherence to information rate using information theory (see Supplementary Material). Unsteady airflow stimulation is inherently unrepeatable so we used the most conservative lower bound method(31) to estimate the information rate for each bin (Fig. 4D). Bins with the highest information rates normalized by spike count are shown for each recorded unit within an animal for both displacement and strain measurements (Fig. 4E). Regions of maximum information rate are broadly consistent for multiple individuals (Fig. 4F), corresponding to the same areas of the wing that our FEA modelling identified as experiencing the largest strains from bend, twist, and camber (Fig. 4G). Overlaying these regions onto a wing sensor distribution map suggests individual CS candidates for the units recorded (Fig. 4H). The highest sensor densities are found where there is the highest information transfer between strain measurement and sensor response (Fig. S7). Sensor activity also remains when the distal portion of the wing is removed (Fig. S8).These are particularly effective locations for observing the strains associated with key wing deformation modes, and away from the wing margins where damage occurs throughout the animal’s life (51). The information rate of dragonfly wing CS is broadly comparable with other spiking sensory systems (Fig. 4I). This is expected, as all mechanosensors abide by the same biophysical principles.

## Discussion

Here, we reaffirm that insect wings, in common with all structures, have finite spatial patterns and ranges of deformation that are excited at particular frequencies. The naturally occurring range of spatial states can be simplified by dimensional reduction methods and are dominated by bending and twisting. When wings are discretized to finite elements with sufficient geometric fidelity, simulations reproduce the same patterns as their primary modes. This FEA model therefore enables detailed analyses of the effect of architectural features and their relationship to sensor distribution (Fig. 4). While bend, twist, and camber may be mechanically inevitable responses for any flexible thin blade, the specific 3D morphology of dragonfly wings shapes the unique bend, twist and camber lines. Wing veins, corrugations, and material heterogeneities collectively enhance and concentrate deformations to specific locations. Placing sensors within these regions of high strain enables more precise tuning and efficient detection of wing loading. Thus, although the presence of these modes (bend, twist, and camber) may be a general mechanical property of flexible wings, our data shows how the dragonfly wing’s detailed architecture facilitates effective sensing and control. Sparse sensors capturing wing deformation is a well-established principle, but our results demonstrate exactly how the dragonfly’s 3D wing geometry shapes the strain patterns and sensor placement on a real wing blade.

Mode-specific strain detection has been observed in other systems. Fly halteres (modified hindwings that detect inertial forces) also demonstrate how biomechanical constraints can shape sensor placement and specificity (10, 13, 14, 18, 48, 52). Whereas haltere mechanosensors are arranged to detect Coriolis forces arising from body rotations, wing mechanosensors also capture the fluid-structure interactions that shape aeroelastic deformations which underly force production and flight control. This is true of all insect wings, where specific morphologies and sensor distributions have arisen as a result of individual behavioural, ecological, and control demands. In dragonflies, we found that the wing’s structural properties act as a computational device that prescribes and constrains deformations into predictable oscillatory patterns, reducing the effective stimulus space prior to neurophysiological transduction. The amplitude of these deformations changes appreciably during free-flight manoeuvres (Fig. 1) and these changes are reflected in the timing of the recorded sensors’ activation (Fig. 3). Comparing the timing of the sensors between wings or relative to the wingbeat cycle would allow dragonflies to monitor the instantaneous loading conditions during these manoeuvres. The timing of these units is not as precise as some other systems, such as dipteran halteres (Fig. S4B; 49), but the decreased timing precision might be explained by dragonfly wings having slower dynamics than fly halteres, more degrees of freedom, and being exposed to less predictable aerodynamic forces in addition to inertial forces. Despite the added complexity of the wing’s stimulus space, we show that these sensors are nevertheless capable of monitoring the natural range of deformations using a simple phase-dependent encoding strategy.

Strain sensors (CS) are most densely clustered in a region of the wing found proximally along the major wing veins where aeroelastic loads give peak strains during gliding and flapping flight. Moreover, this evolved sensor placement was predicted by the areas offering the highest information rates for the recorded units. There is strong spatial correlation across the wing, so sensors could potentially be placed in many different locations and extract similar information. However, small differences in sensor accuracy can have a huge impact when monitoring complex behaviours like flight, and it is unlikely that these sensors are placed by chance in the region which contains the most accurate information. Prior work on generic deformable wings produced optimal sensor placements and information patterns that differ from what we observed in dragonflies (28, 29). These locations were influenced by wing stiffness and neural threshold, but in general, modelled sensors tended to cluster along the outer edges of the wing tip or base. In contrast, insect wing strain sensors are most often concentrated at the wing base along the interior veins (4, 6, 53). This comparison supports the idea that natural wing morphology specifically shapes sensor placement and information encoding in ways that cannot be captured by simplified flat wing models.

Dragonflies have hundreds of wing mechanosensors(5), yet we found just a few large amplitude units are activated and dominate the signal in the wing nerve under natural conditions (in the absence of perturbation). The additional sensors recruited during white noise perturbations may be necessary under circumstances where unusual temporal and/or spatial deformations occur and are likely tuned to higher velocity changes, consistent with the fact that white noise perturbations contain higher frequency displacements than airflow-induced flutter. Our results suggest that while the wing sensory system uses few sensors to monitor the wing state, many additional sensors are present and poised to detect deviations from the normal expected modes – i.e. continuous oscillations primarily composed of bend and twist. The speed at which dragonflies must react to changes in their expected sensory environments and the number of sensors recruited during these perturbations indicates that deviations from the natural wing state are likely represented as a population code. This population code may also be adaptable. Sensor locations and even axonal paths can vary between individuals(5) and dragonflies can experience significant wing damage throughout their lives(51). We hypothesize that when wing mechanics change – either during development as they harden over time, or if they are damaged - adaptation is accomplished further downstream by integrating interneurons that receive inputs from multiple modalities. In this case, visual, ocellar, or proprioceptive information would also be necessary to augment the wing sensory inputs when the wing mechanics change. Other senses could provide a ground truth comparison for the animal’s perception of self-motion which would allow recalibration of their sensorimotor control internal models(54, 55).

Invertebrates have historically provided a significant source for bioinspiration in engineered technologies(56–62). There have been many studies on dragonfly wing aerodynamics and some on material properties, yet 3D anatomy has rarely been addressed with sufficiently high fidelity for accurate structural analysis and assessed in the context of free-flight. Similarly, the distribution of mechanosensor types have been identified(5) but this is the first characterization of how the loading states are monitored. Accurate state estimation using wing- mounted strain sensors has recently been demonstrated as a promising avenue for the development of ‘fly-by-feel’ biomimetic aerial vehicles(63, 64). Other deformable objects, from bridges to prosthetics, have been extensively studied across fields with much effort placed on displacement and strain field measurement and reconstruction techniques (65–72). Here, we have shown how morphology dictates and limits the range of the stimulus space, and strategically placed sensors enable sufficient sensitivity, while reducing detection of redundant signals. The dragonfly’s morphology and sensor distribution allow their relatively small nervous systems to produce simple and scalable representations of natural deformations.

## Materials and Methods

### Free-flight kinematics

*S. striolatum* were filmed taking off from a vertical perch using nine high-speed cameras (five Photron AX200 and four Photron SA3, Photron Ltd) recording at 2000 fps and 1024 x 1024px. Sequences were trimmed to exclude time when the body was within approximately one body length of the perch and all four wings were engaged in stereotypical flapping flight. A voxel- carving method was used to measure the six degrees of freedom of the body and wingtip spherical coordinates(73). Fifth order polynomials were fitted to the voxels corresponding to the leading and trailing edge of the wing (from 10% to 90% wing length), which were then transformed into the wingbase reference frame, aligned with the basal twist of the wing. Wing deformations (MovieS4; Fig. 2Di) were then calculated by fitting a 80x50 point surface to the leading and trailing edge polynomials. In total, 14 sequences were processed, including 99 wingbeats (pooling left and right forewings).

### Nerve ablations

*S. striolatum* were collected from artificial ponds at Imperial College London Silwood Park campus. Animals were kept at 7°C prior to experiments and restrained on a 13°C Peltier during surgery. Animals were subjected to either a sham surgery where the anterior nerves were exposed but kept intact or an ablation surgery where the anterior nerves were lesioned. After surgery, parafilm was used to cover the incision site and secured with beeswax. Animals were allowed 20-30 minutes recovery time and were screened prior to and after surgery to ensure they were healthy and capable of sustained lift in free-flight. This was defined by whether, when dropped, individuals were able to navigate to the ceiling or wall of the filming tent at or above the initial drop height. To encourage animals to take off, they were placed on a 75x75 mm platform with a Peltier heating element underneath. The Peltier warmed to a maximum of 59°C over 1.5 minutes. Take-offs were filmed at 1000 fps using two Chronos 2.1-HD cameras (Kron Technologies Inc., Canada) and digitised using DLT DataViewer(74). Data were not normally distributed, so comparisons were made using Kruskal-Wallice (WBF) or Multi-sample test for equal median directions (phase) with Bonferroni correction for six comparisons (Table S1)(50).

### Deformations

To identify the global motion patterns of the wing we aimed to represent the overall wing state with few variables. To measure local motion and strain, we filmed the fluttering wing during electrophysiological recordings at 1000 fps using a Chronos 2.1-HD camera (Kron Technologies Inc., Canada) and performed digital image correlation (DIC)(75) on the recorded video frames (Fig. 2B). DIC requires a material’s surface to be patterned, typically by painting with a splatter coat, however painting the wing would alter its mechanics and we found that the wing venation pattern alone provides sufficient visual patterns for DIC to extract displacement and strain. To accommodate electrode and stimulus while maximising visualisation of the largest deflections (dorso-ventral), animals were positioned on their side and angled upward at 40° with the camera placed approx. 20° behind the animal. Perspective correction, cropping, and contrast enhancement were performed on each frame prior to DIC analysis. Subset window sizes varied slightly between animals depending on the wing size and amount of motion blur but were typically 1% of the total wing size. Single camera (2D-DIC) measurements with perspective correction were sufficient to capture the same measured displacements as two camera (3D-DIC) measurements(76) (Figure S3A).

### Electrophysiology

The wing was stimulated using either airflow or direct mechanical vibration. The airflow stimulation apparatus consists of an axial fan and airflow straightener to blow air across the wing at behaviourally relevant speeds (1-3m/s) and angles of attack (-/+20° & 0° relative to the chordwise plane between the costa and arculus veins; Fig. S3). The direct mechanical stimulation was provided by a precision lever-arm system (Aurora Scientific 300C: Dual-Mode Muscle Lever). The lever was attached at the nodus on the wing, and vibration stimuli included uniform white noise (0-500Hz) or aeroelastic flutter playback (Fig. S5). The flutter displacement data were derived from digitising the nodus during airflow stimulation using DLT DataViewer (74). This stimulus replicated the natural nodus motion in the absence of airflow. The forewing was fixed at the hinge in a natural (gliding) flight position using a 50/50 mixture of paraffin and beeswax.

Fixing the hinge largely excludes internal proprioceptors from stimulation, allowing only wing surface sensors to be activated. The airflow stimulation apparatus or lever-arm system was aligned to the fixed wing sample rather than to the body axis to account for any difference across specimens. Hindwings were folded dorsally out of view of the camera and restricted with a folded strip of paper and paper clip. To record the activity of wing mechanosensors, we performed a side entry dissection and removed a single layer of trachea to access the anterior wing nerve. A borosilicate glass suction electrode was pulled using a Sutter P-2000 Laser-Based Micropipette Puller, cut with a ceramic tile to match the diameter of each nerve (30-50 μm), and used to record extracellularly from anterior nerve primary afferents. Anterior nerves receive inputs from all wing vein CS (Fig. 4H). The posterior wing nerve only receives inputs from the wing hinge, so posterior nerve recordings were not included here. Recordings where muscle contractions occurred were easily identified from the large movement artefacts and were subsequently discarded.

### Finite Element Analysis and Fluid-Structure Simulation

The wing’s structural properties largely dictate the modes of deformation. Combes and Daniel (2003) found that the hawkmoth’s wing deformation patterns are still sufficiently predicted when fluid-dynamic forces are excluded from damped finite element models(77, 78). Thus, we also performed a modal analysis simulation for a dragonfly forewing model retaining realistic corrugation with Ansys Mechanical 2024 R2 (Movie S5). The wing model geometry was constructed by combining the µCT scanning for a *S. striolatum* wing for vein cross-sections and the 3D venation from the direct linear transform (DLT) from a pair of stereo photos. See Fabian et al., 2022 for the details of the model construction method. The wing density (except for the pterostigma) was assumed to be 1200 kg/m3, while the pterostigma mass was set to be 9% of the total wing mass(79). The Young’s modulus was 4.9 GPa for the entire wing(80). Using the same wing model, we performed a fluid-structure interaction simulation on a gliding dragonfly forewing. The fluid dynamics solver (Ansys Fluent) for unsteady laminar simulation and structural dynamics solver (Ansys Mechanical) for transient structural simulation were loosely coupled (Ansys System Coupling) to obtain the converged solution for each time step. The example sequence shown in Movie S1 is a gliding simulation where wind speed is 2.6 m/s and angle of attack is 15° (Reynolds number is approximately 970).

### Temporal Analysis

In this study, STA was sufficient in providing the first approximation to the sensory encoding especially when the input information was linearly represented. Non-linear models such as maximum noise entropy (MNE) and maximally informative dimensions (MID) were not applicable for several reasons. Firstly, the recorded sensors are sparsely activated, and the recording cannot be maintained for long periods due to their proximity to the mechanical stimulation. The limited spikes led to overfitting in MNE and MID with uninterpretable results.

Secondly, the stimulus space dimensionality of a fast deforming wing is an order of magnitude larger than what is typically used for non-linear models(81). To accurately represent the local deformations across the wing surface requires ∼30,000 variables (bins) per time point (frame). Typically, only 1-2 variables are used in these models, so the computational requirements for 30,000 are not realistic. To reduce computational requirements, dimensionality reduction techniques can be performed prior to modelling, but this requires making more assumptions about the data. Units which were not phase-locked with the periodic stimuli were excluded from STA analysis (see Fig. S4B for further explanation). Phase-locked units were classified using vector strength (Fig. S4)(49, 50). Phase was calculated for each unit relative to PC scores by measuring the time between oscillation peaks and normalizing this to radians. The time point between peaks where the spike occurs is the phase value in radians. Phase-locked units were identified by calculating vector strength (see Eq.1). Units with vector strength >0.6 were considered phase-locked and vector strength between 0.5-0.6 were classified as weakly phase-locked. Units that were not phase-locked were likely a result of inaccurate sorting due to the low signal to noise ratio of smaller units.

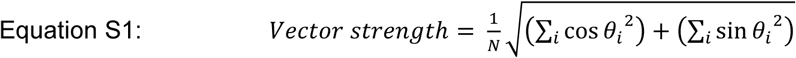

### Perturbations

In addition to white noise vibrations, we also replayed natural deformations generated from airflow-induced oscillations. Airflow-induced oscillations and airflow playback vibration PCs are the same, but with different rank orders (Fig. S5A). Airflow playback resulted in an intermediate sensory response with more recruitment than airflow, but less than white noise vibration (Fig. S5B). These differences were likely a result of the frequency content (Fig. S6) of vibration-induced deformations being different from airflow-induced and free-flight. The amplitudes tested were initially matched to airflow-induced oscillations. However, smaller amplitudes activated many more units for vibration-driven experiments, so while a similar range of amplitudes were covered, on average, lower amplitudes were tested for perturbation experiments. The mean displacement from baseline (in either direction) for airflow-induced deformations was 2.6 +/- 1.2 mm and the maximum displacement tested was 13.6 mm. The mean displacement from baseline (in either direction) for vibration-induced deformations was 1.2 +/- 1.1 mm and the maximum displacement tested was 15.4 mm. The maximum (positive direction) displacement measured across trials for free-flight deformations was 2.5 +/- 0.5 mm and the minimum (negative direction) was 2.1 +/- 0.5 mm.

### Neural information

The neural response variable x represents the mechanosensory spike train converted to a binary vector at the video frame rate (1 = spike, 0 = no spike), aligned to video frames and mean subtracted prior to coherence analysis. This approach preserves spike timing at the video frame resolution without additional smoothing. The magnitude-squared coherence (C_xy_) was estimated between each unit’s activity (x) and the displacement or strain change from baseline of each bin (y) as a function of their power spectral densities, Pxx and Pyy, and cross power spectral density, Pxy between 0 and 500Hz(82). Numerical approximation error was reduced by increasing the number of segments (dividing the input signals into smaller, overlapping 100 ms sections). This allows for better averaging and reduces the influence of random fluctuations within individual sections. Averaging across many segments decreases the frequency resolution, which is reflected in Figure 4A where the frequencies measured appear smoothed. This can also result in a loss of information for slowly varying components of the signal; however, these components are not behaviourally relevant as the forces acting on the wing during flight are not slowly varying. Coherence calculations using few segments did produce similar spatial maps but also amplified DIC measurement errors resulting in unreliable coherence and information rate estimates.

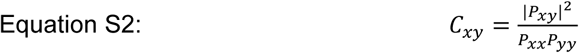

Airflow stimulation is generally only statistically repeatable, so the more conservative lower bound method was used to estimate information rate (I) in bits/s from coherence, which assumes linearity and stationarity. (31, 83). For most analyses, we used displacement as a more direct representation of the wing movement. However, for the sensor localization (Fig. 4), strain was the more appropriate choice as the local strain pattern is what activates these sensors.

Occasially, DIC analysis could not resolve wing edges due to motion blur, so those pixels were not included in coherence estimations. We did not remove entire frames, only the individual pixels that could not be resolved. These pixels tended only to occur along the lateral trailing edge of the wing where sensor density is far lower. To normalize for spike rate, high information regions were defined as the bins with information rates within the top 1-5% (depending on the number of unresolvable pixels removed) for each individual unit.

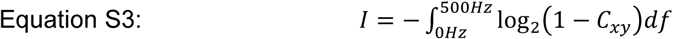

The mean information rate for dragonfly wing mechanosensors was found to be higher than other systems (Fig. 4I), in part because we used turbulent airflow instead of white noise (which covers a larger range and more even distribution of frequencies).

### Imaging

See Fabian et al., 2022 for neural imaging and sensor distribution mapping protocols.

## Acknowledgments

The authors thank Daniel Ko, Sean Lim, Janka Kluga, Rui Zhou, and Myriam Uhrhan for their help with instrumentation. We thank Holger Krapp, Benjamin Campbell, and Samuel Fabian for their valuable discussion and feedback.

## Data availability

Data are available at Dryad Digital Repository: DOI: 10.5061/dryad.41ns1rnqh

## Author Contributions

Conceptualization: AMY, HTL, RJB; Methodology: AMY, HK MM, SMW, RJB, HTL; Investigation: AMY, HK MM, SMW, RJB, HTL Visualization AMY, HK MM, SMW, RJB, HTL; Supervision: HTL, RJB; Writing: AMY, RJB, HTL

## Supporting Information Text

### Model choice

To demonstrate how morphological computation contributes to the tuning of biological sensory systems, dragonflies are an ideal model. Their large, highly deformable wings are easy for structural measurements and their wing afferent signals are accessible for electrophysiological recordings. Furthermore, while all flying animals have deformable mechanosensitive wings, dragonflies are among the most agile fliers and their wings are frequently used as models for aerodynamics and wing mechanics(37, 76–84). The dragonfly wings’ mechanics naturally limit their range of deformation, but there is still a large available state-space significant for flight control.

### Tegula complex

In locust(1, 2) and hawkmoth(3) a mechanosensory structure on the hindwing - called the tegula - influences forewing motor control. The tegula plays an important role in the generation of rhythmic flight motor patterns in both insects but has not previously been described in dragonflies. We identified a tegula-like structure called the ‘tegula complex’ which comprises a hair plate and CS field (Fig. S8D). The tegula complex, like the tegula, may be responsible for maintaining wing synchrony in dragonflies. Forewing nerve ablated dragonflies are still able to maintain synchrony between their fore- and hindwings, albeit at a lower frequency, indicating that the hindwings (and hindwing tegula complex) may be responsible for maintaining wing synchrony. Afferents from the tegula complex were not recorded here but may be of interest for future studies investigating mechanisms for wing synchronization.

### Phase shifts

Phase time shifts that result from changes in displacement amplitude (Fig. 3C) are descriptive of a single feature only and those features have been extracted from the overall displacement pattern. Larger amplitude bending does not scale equally with twist or camber, and here the displacement speed has also been normalized across amplitudes. This may partially explain why we often see a phase advance rather than a phase delay with increased amplitudes. Alternatively, large amplitude bending is typically caused by large aerodynamic loading. Such passive deformation can be characterized by a wave propagating tip-to-base, in contrast to the deformation driven by the flapping musculature with a base-to-tip propagation. Depending on the location of the exact strain sensor being recorded, the phase timing will shift according to the contribution from these two waves. Regardless of the specific physical mechanism, it is important to consider that the overall pattern of motion is not identical for every amplitude. However, because the stimulus space is constrained by the wing structure, the possible deformations are still limited to a finite range of patterns that can be described by bend, twist, and camber. Any motion outside of the natural wing state will necessarily activate additional sensors that can encode the deformations not represented within the natural range.

### Perturbation-activated sensors

In general, individual variation and adaptability exist because the nervous systems is able to produce appropriate outputs using a variety of different mechanisms(4, 5). We did not characterize the selectivity of perturbation-activated sensors as it is outside the scope of the current study. It requires extensive cataloguing of responses across conditions, morphologies, and individuals. Selectivity and encoding strategies are unlikely to be identical between individuals(5) and input sensitivity is also likely to depend on context e.g. structural changes or behaviour state(6–8). The large number of active units also prevents accurate sorting with the current recording preparation. Future experiments will aim to resolve more units simultaneously for characterizing these sensors’ selectivity and other factors including sensor redundancy and adaptability.

### Deformation Frequency components

Airflow induces consistent wing oscillations as a result of the structural and material properties of the wing. The natural frequency of the wing (Fourier transform dominant frequency of mean displacement of entire wing) across animals was 135+/-23Hz. Faster air speeds resulted in slightly higher frequencies (18.6+/-15.2Hz increase from 1-3m/s, N=5 animals). The dominant frequencies of the PCs were the same as the mean displacement frequency (Fig. S6). Vibration induced deformations have higher frequencies of twist and camber motion.

**Fig. S1.**
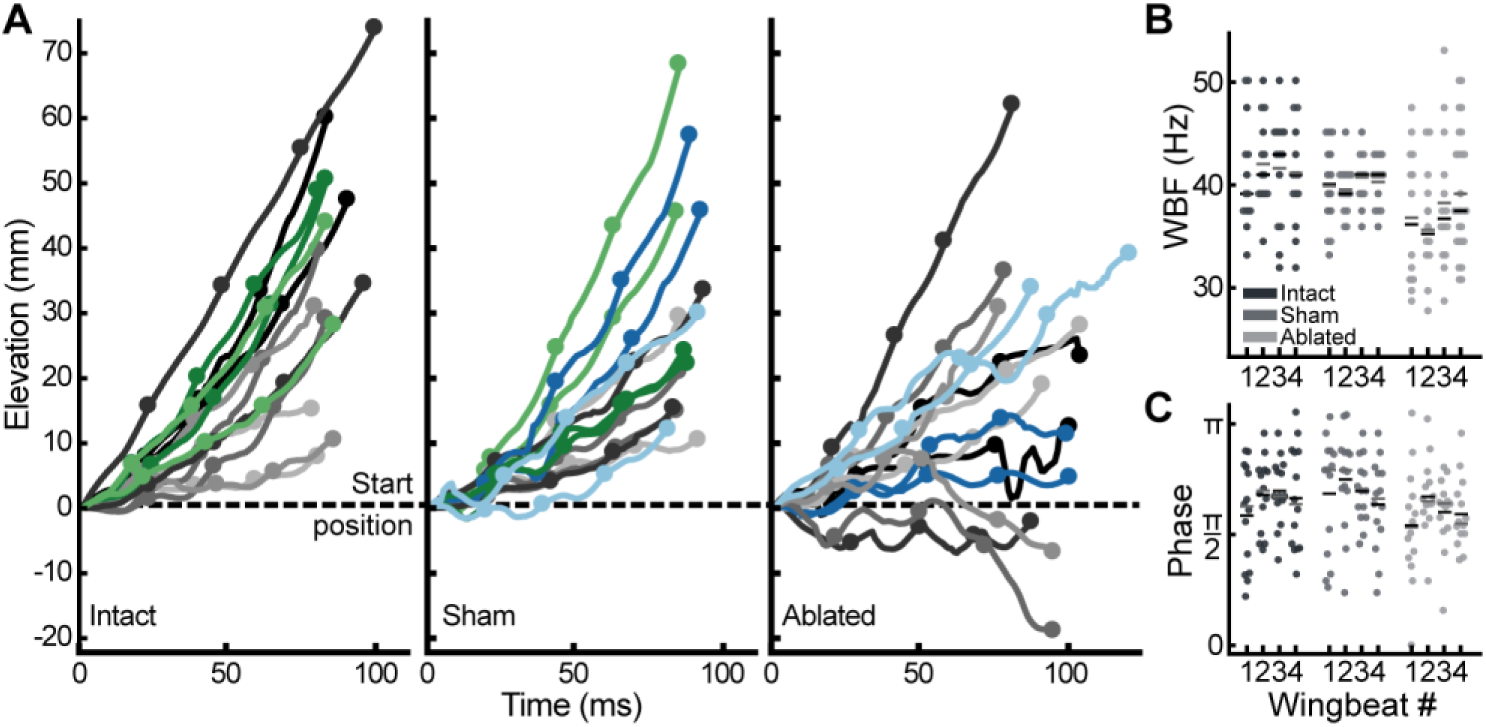
Nerve ablation. A) Take-off elevation over the first four wingbeats. Different shades represent different animals. Circles mark start of forewing downstroke. N=7 animals, 2 trials per condition. Two animals appear in both intact and sham (green) and two in both sham and ablated (blue). B) No significant differences in WBF between wingbeats within each condition. Grey and black bars show mean and median respectively. Ablated WBF significantly lower than intact (P=0.00002) and sham (P=0.0024). No significant difference between intact and sham WBF. C) No significant differences in the hindwing phase relative to the forewing between wingbeats within each condition. No significant differences between all conditions. Data resolution results from video frame rate (1000 fps).

**Fig. S2.**
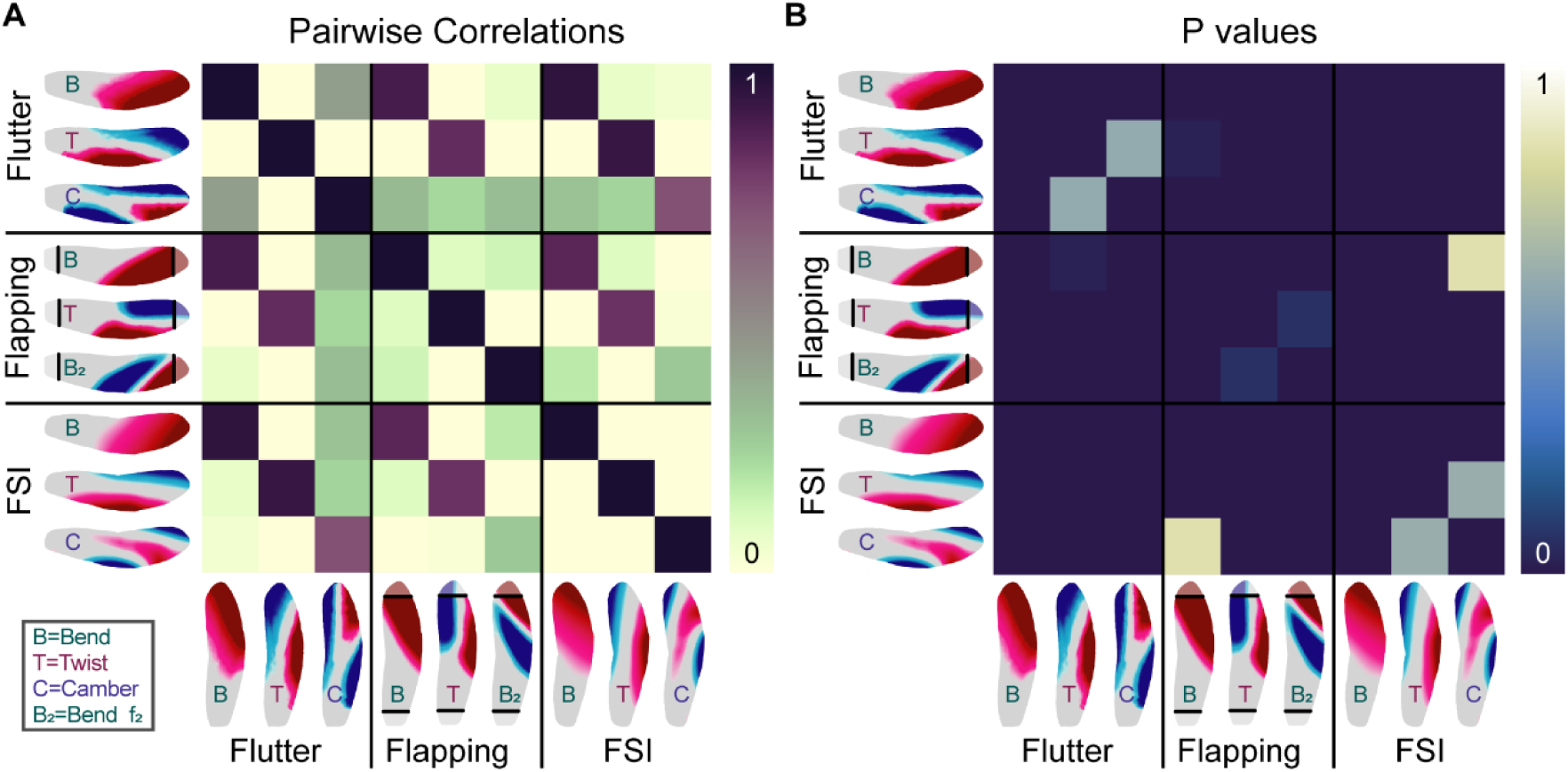
Pairwise correlations of displacement maps. A) The bend pattern is highly correlated across all conditions. Twist is also highly correlated between conditions. Camber is only correlated between Flutter and FSI because camber could not be measured during flapping. B) All highly correlated pairs were significant. The only non-significant correlations were between combinations with low correlation values (e.g. flapping bend & FSI camber).

**Fig S3.**
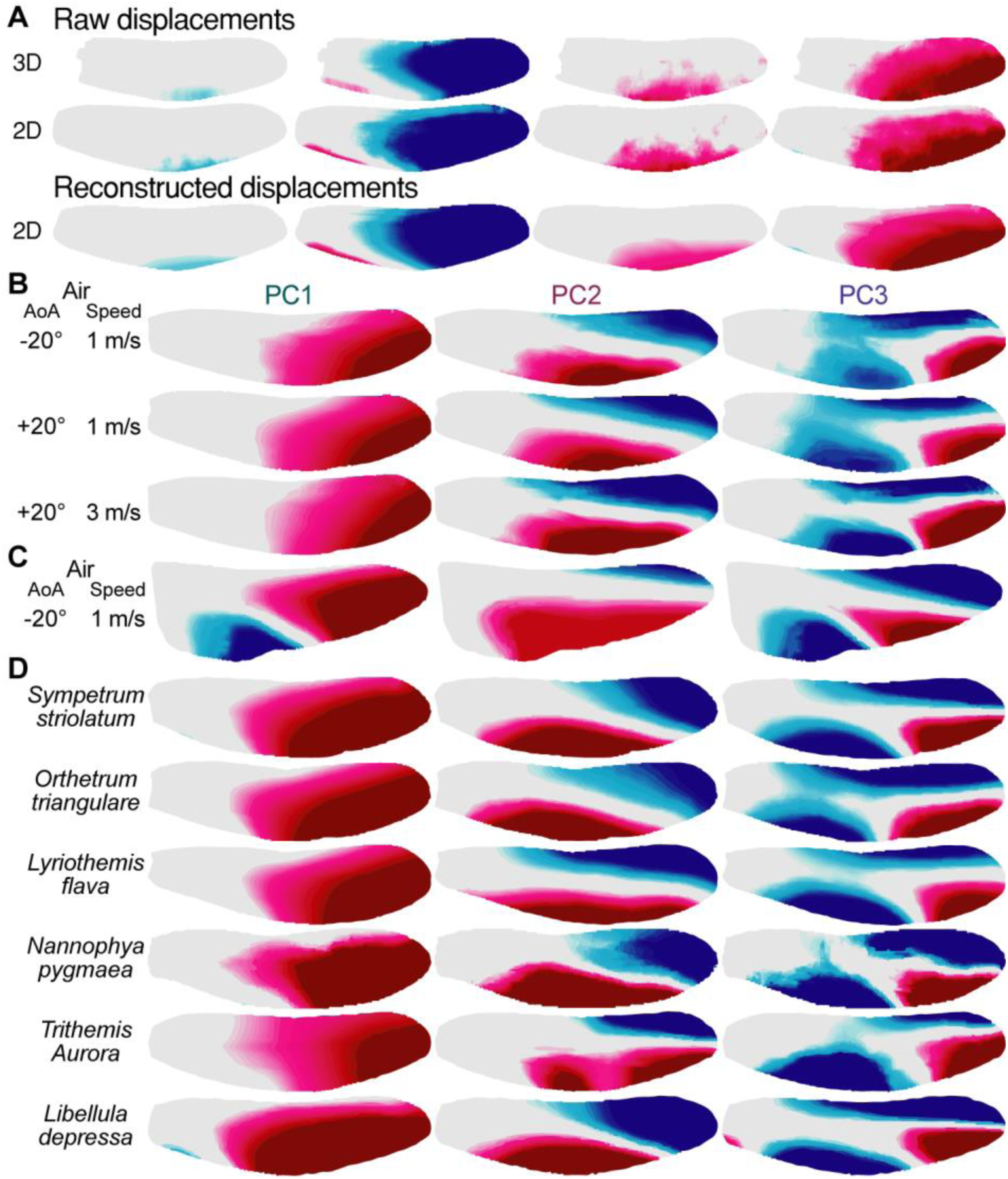
Wing displacement PCs. A) Example frames of two camera 3D measurements of displacement magnitude (top), single camera dorso-ventral displacement measurements (middle) and displacement reconstructed from PCs 1-3 (bottom). B) Angle of attack (AoA) and air speed do not influence PCs. Example PCs calculated from 7000 frames. C) Hindwing PCs are similar to forewing PCs. PCs calculated from 5000 frames. D) PCs are the same across dragonfly species tested. PCs calculated from 100-1000 frames. All wings filmed at 1000 fps and resized to equal wingspan and cord width for easier comparison.

**Fig. S4.**
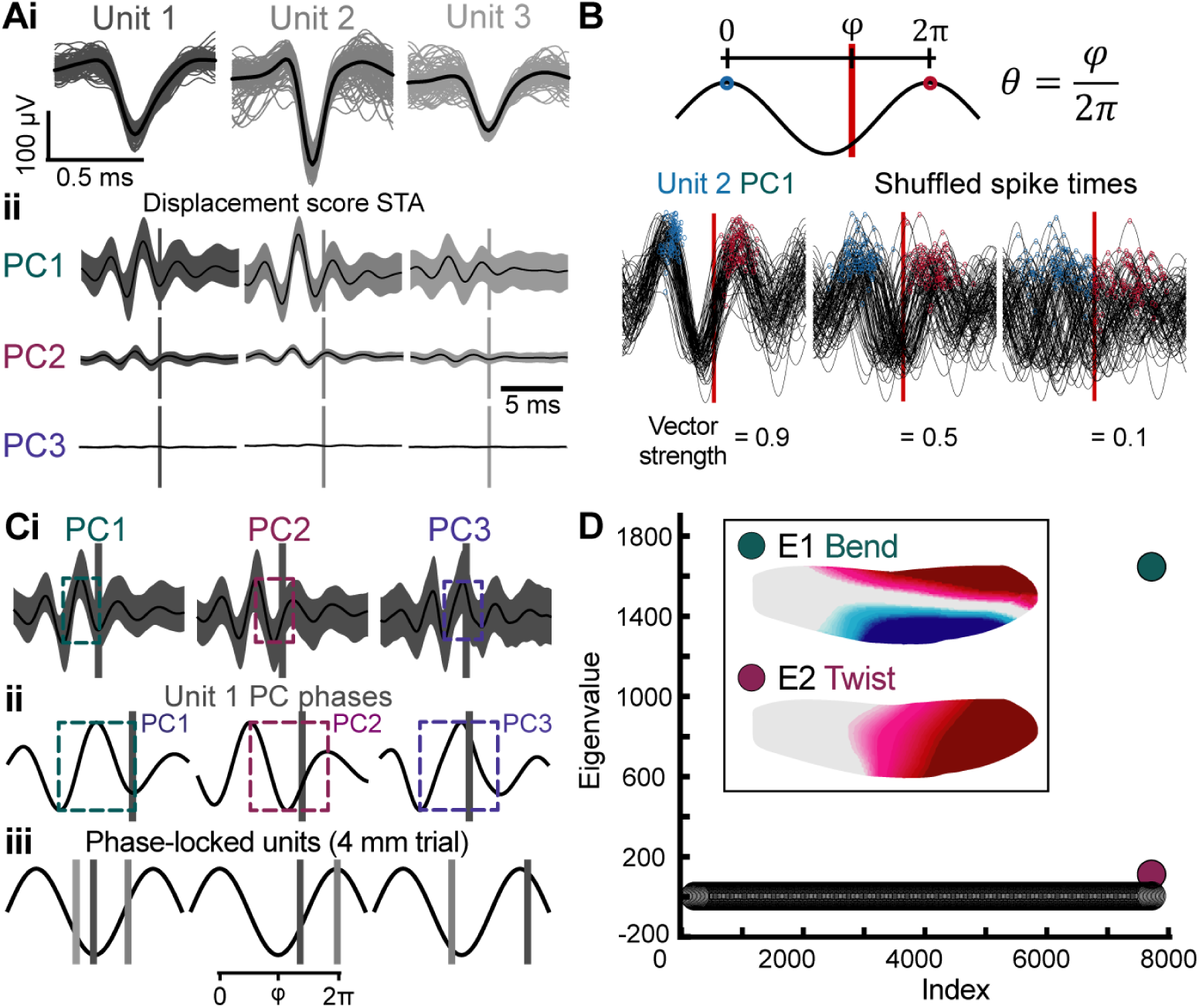
Phase measurements. A) Sorted units shown in Figure 3 (i) and their associated PC STAs (ii). B) Phase (q) is calculated by measuring the relative time (j) between oscillation peaks normalized to radians. Phase-locked units are identified by calculating vector strength (see Eq.1). Units with vector strength above 0.5 are classified as phase-locked. C) Unit 1 phase (vertical bars) relative to PC score STAs (i) normalised to radians (ii). All phase-locked units from airflow induced deformations with 4 mm mean displacement (iii; see also Fig. 3). D) Significant features (E1-E2) identified through covariance analysis of the spike-triggered raw displacements. Camber was notably not present, indicating that this mode is less likely to be represented in the neural signal. <u>This result was consistent across units tested</u>.

**Fig. S5.**
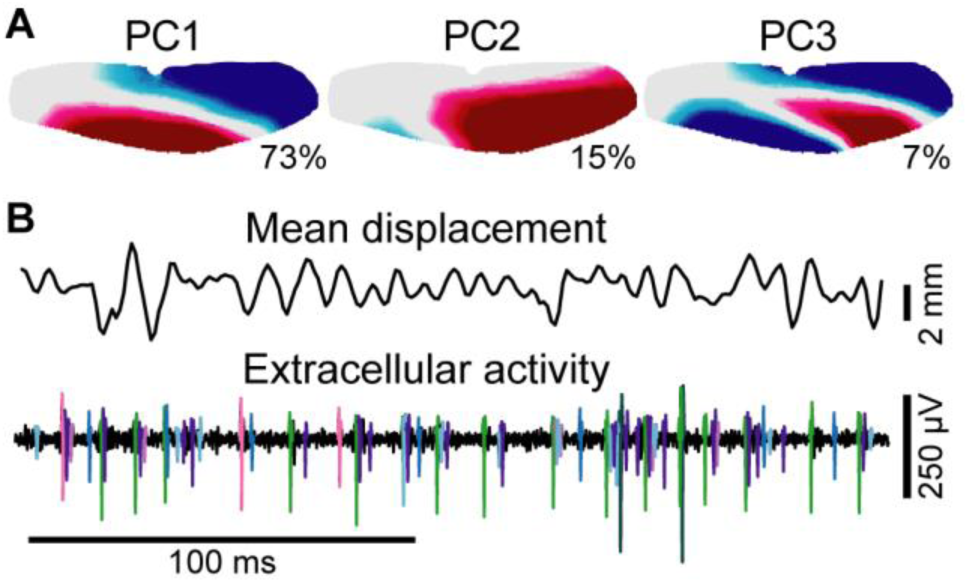
Airflow playback vibration. A) Airflow playback vibration induced deformation PCs. B) Airflow playback result in moderately increased extracellular activity compared to airflow induced displacements. Colours indicate different units.

**Fig. S6.**
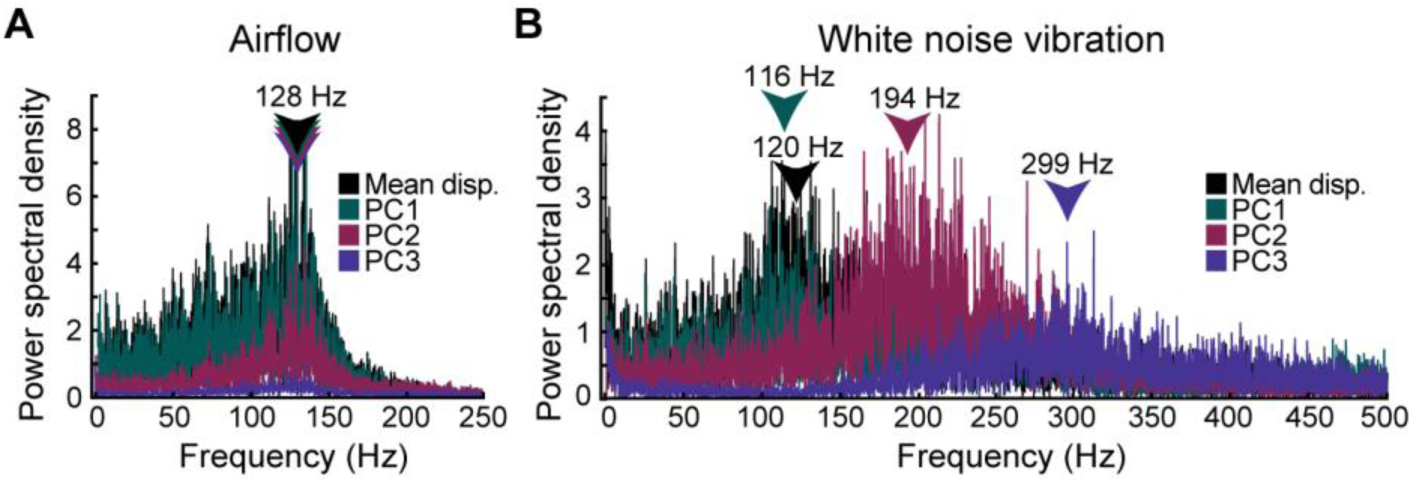
Displacement frequency components. Frequency components of mean absolute displacement of the entire wing and PC scores (PC1-bend, PC2-twist, PC3-camber) for airflow (A) and white noise vibration (B) stimuli.

**Fig. S7.**
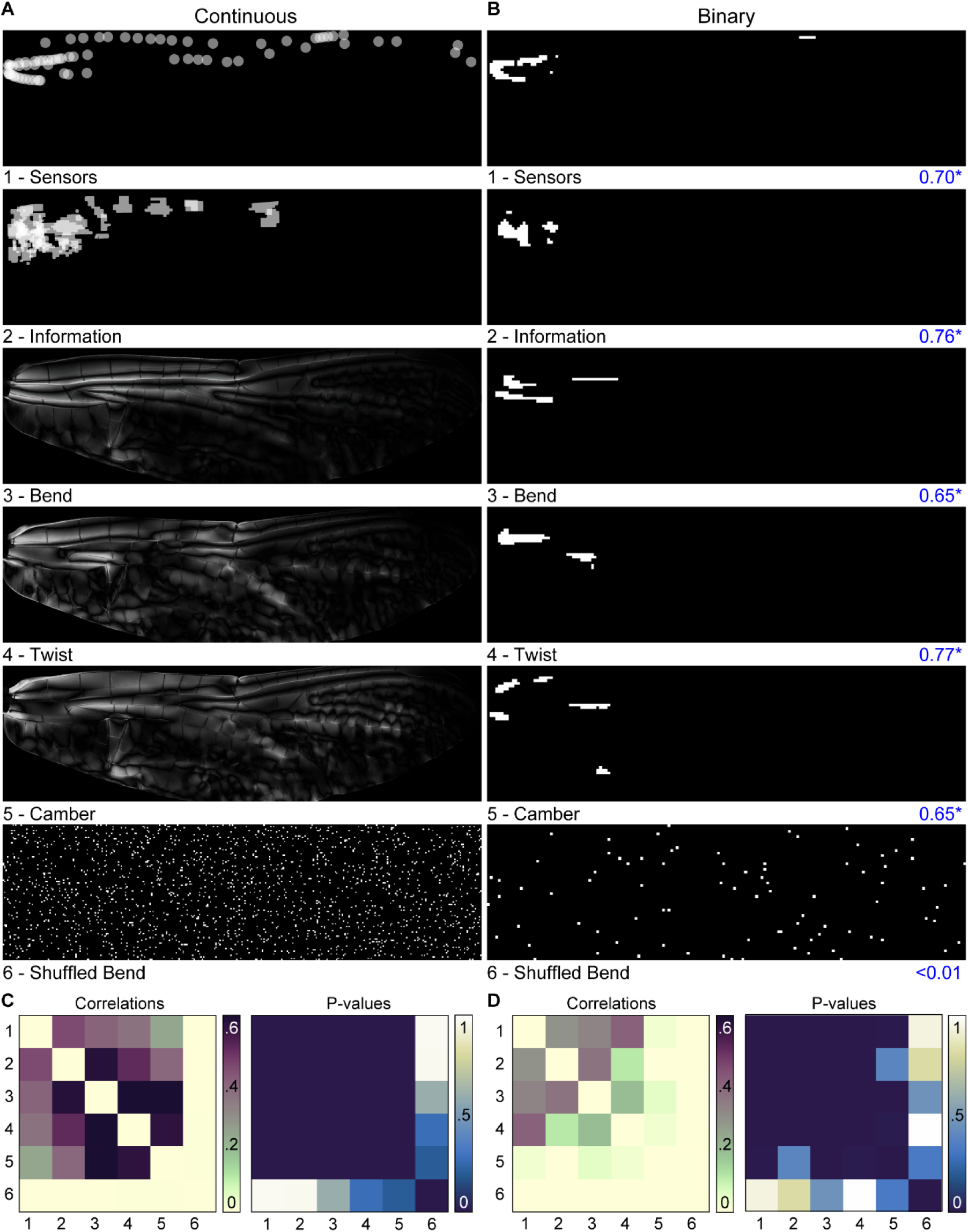
Hotspot analysis. A) Sensor, information, strain maps, and shuffled bend stain map converted to continuous greyscale images. B) Hotspots from A defined by 99^th^ percentile plotted as binary images. Spatial autocorrelation (Moran’s I) values for each image shown in blue. Values >0.5 suggest strong clustering. Significant values indicated by *. C) Correlation statistics for continuous images (A). D) Correlation statistics for binary images (B).

**Fig. S8.**
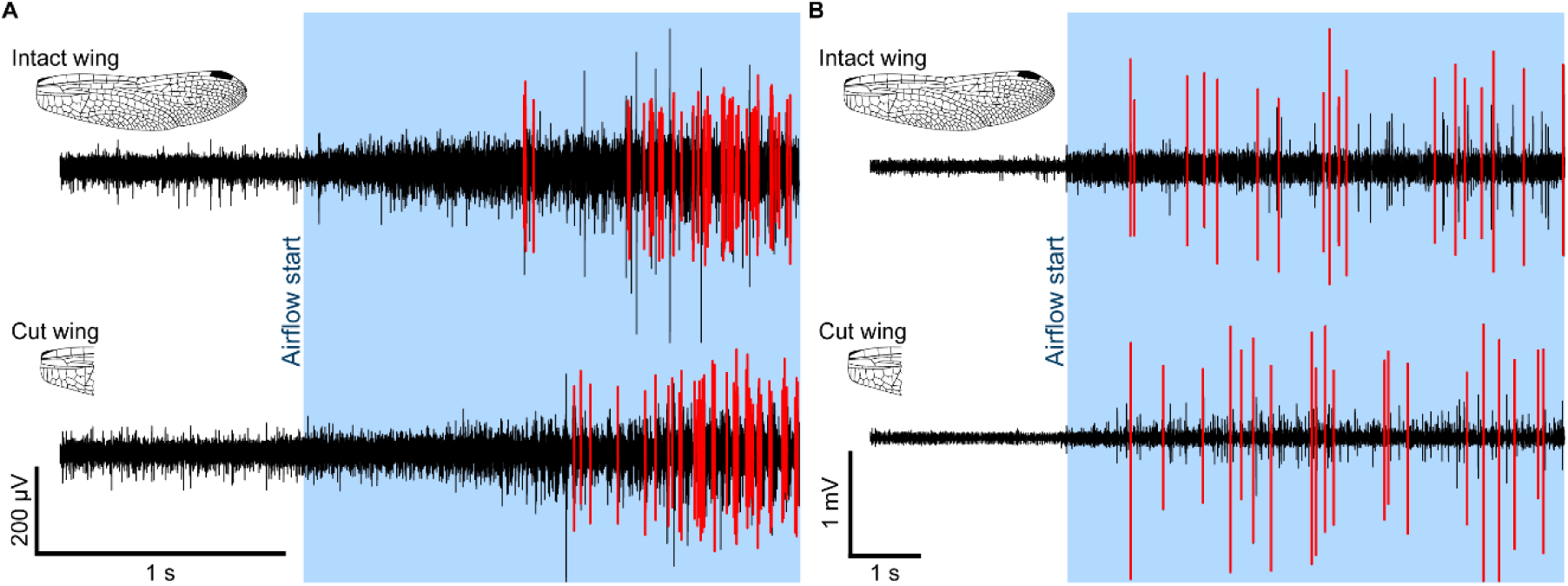
Sensor activity remains after distal portion of wing is removed. Extracellular recording from *Orthetrum cancellatum* (A) and *Celithemis elisa* (B) wing nerve in response to airflow with wing intact (top) and cut (bottom). Large sensory units highlighted in red.

**Fig. S9.**
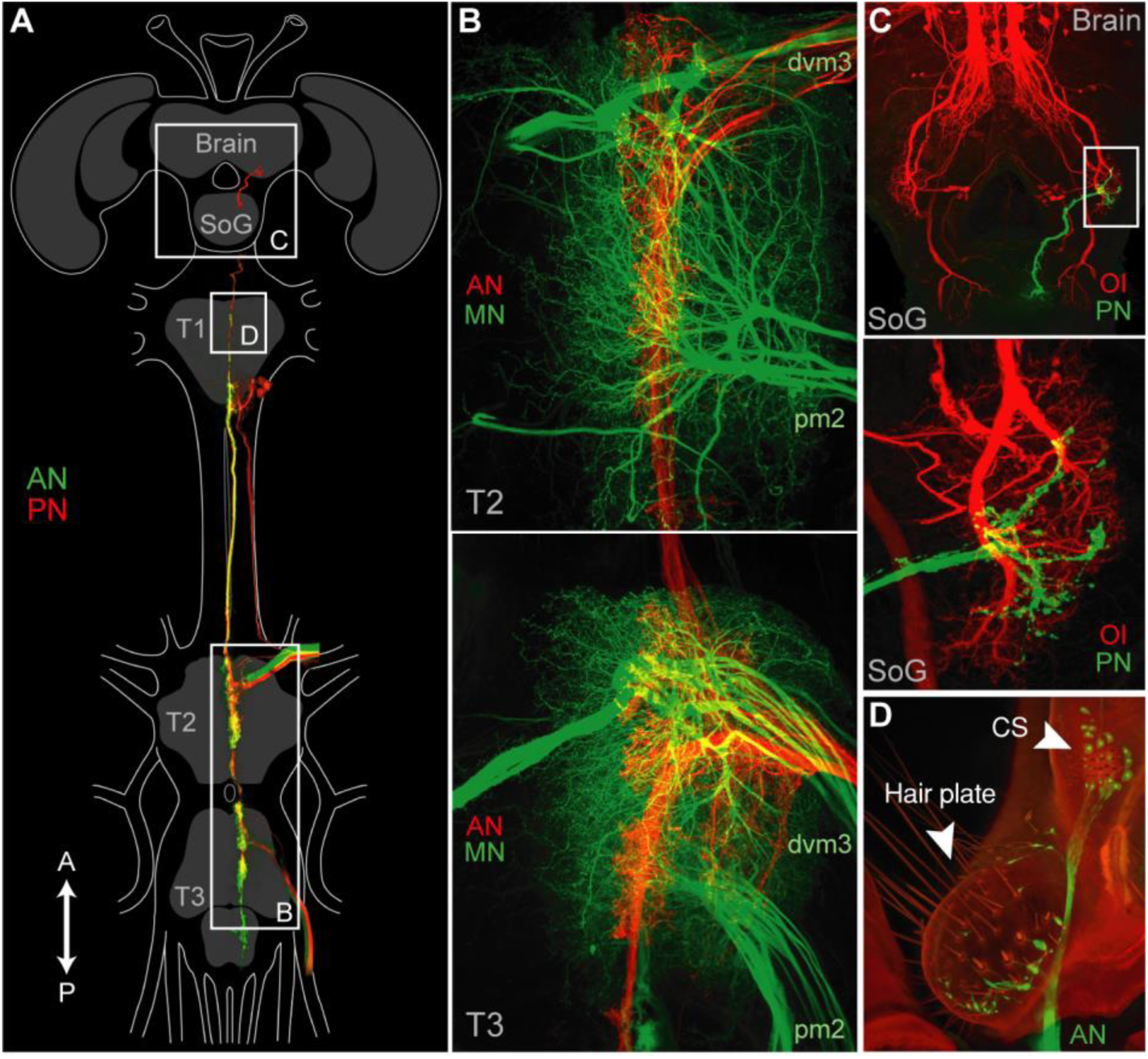
Wing mechanosensor neuroanatomy. A) Anterior (AN) and posterior (PN) wing nerves project ipsilaterally within T2 & T3. PN projects to T1 and brain (panel C) and a subset of AN afferents from the tegula complex (panel D) project to T1. B) AN projects to wing motor neurons (MN) dvm3 (pronation) and pm2 (anti-overpronation). C) PN projections to secondary ocellar interneurons (OI) in the posterior deutocerebrum. D) Hair plate and CS comprise the tegula complex.

**Table S1.**

| Comparison | Between wingbeats |  |  | Between conditions |  |  |
| --- | --- | --- | --- | --- | --- | --- |
|  | Intact | Sham | Ablate | Intact vs. sham | Sham vs. ablate | Intact vs. ablate |
| Phase | 0.58 | 0.47 | 0.75 | 0.058 | 0.023 | 0.57 |
| WBF | 0.94 | 0.63 | 0.49 | 0.016 | 0.0024* | 0.000017** |

Multiple comparisons P values for nerve ablation experiments.

---

**Movie S1.**

High-speed video of a dragonfly taking off in its natural environment followed by the same behaviour captured in a nine-camera, free-flight arena. Wing kinematics of freely flying dragonflies were tracked (left) and reconstructed in 3D (right). Wing displacements (blue-upward, red-downward) were measured throughout the stroke cycle during free-flight and simulated gliding (fixed-wing airflow-induced displacements). Wing displacements were measured while simultaneously recording the neural activity of wing mechanosensory afferents (strain sensors). 99% of the measured displacements are comprised of three principal components (bend, twist, and camber) and different sensors (coloured spikes) are activated by different ratios of those components. Micro-CT scans of the wing were used to construct a high-fidelity morphological model of the dragonfly forewing. Fluid-structure simulations of gliding flight show detailed measurements of wing displacement and strain. The regions of the wing that experience the largest strains overlap with the identified locations of the strain sensors (that encode the principal deformation components).

**Movie S2.**

Free-flight kinematics: Example flight trajectory includes three wingbeats where a roll manoeuvre is performed during the last wingbeat.

**Movie S3.**

Example intact vs. nerve ablated take-off. Wingbeat frequency decreases after nerve ablation and animals often abandon take-off attempts once airborne.

**Movie S4.**

Free-flight deformations. Displacement field maps generated from tracked leading and trailing wing edges.

**Movie S5.**

Global displacement of FEA modes. Modes 1 & 2 show bend and twist. Modes 3-4 show camber.

## References

1. S. Lee, J. Kim, H. Park, P. G. Jabłoński, H. Choi, The Function of the Alula in Avian Flight. Sci Rep 5, 9914 (2015).

2. R. Wootton, The Geometry and Mechanics of Insect Wing Deformations in Flight: A Modelling Approach. Insects 11, 446 (2020).

3. R. Wootton, Support and deformability in insect wings. Journal of Zoology 193, 447–468 (2009).

4. B. R. Aiello, et al., Spatial distribution of campaniform sensilla mechanosensors on wings: form, function, and phylogeny. Current Opinion in Insect Science 48, 8–17 (2021).

5. J. Fabian, et al., Systematic characterization of wing mechanosensors that monitor airflow and wing deformations. iScience 25, 104150 (2022).

6. B. Pratt, T. Deora, T. Mohren, T. Daniel, Neural evidence supports a dual sensory-motor role for insect wings. Proceedings of the Royal Society B: Biological Sciences 284, 20170969 (2017).

7. D. C. O’Carroll, N. J. Bidweii, S. B. Laughlin, E. J. Warrant, Insect motion detectors matched to visual ecology. Nature 382, 63–66 (1996).

8. H. C. Bennet-Clark, “8. Songs and the Physics of Sound Production” in *8.* Songs and the Physics of Sound Production, (Cornell University Press, 2019), pp. 227–261.

9. K. Schildberger, M. Hörner, The function of auditory neurons in cricket phonotaxis. J. Comp. Physiol. 163, 621–631 (1988).

10. A. Fayyazuddin, M. H. Dickinson, Haltere afferents provide direct, electrotonic input to a steering motor neuron in the blowfly, Calliphora. J Neurosci 16, 5225–5232 (1996).

11. B. Webb, R. R. Harrison, M. A. Willis, Sensorimotor control of navigation in arthropod and artificial systems. Arthropod Structure & Development 33, 301–329 (2004).

12. K. D. Longden, T. Muzzu, D. J. Cook, S. R. Schultz, H. G. Krapp, Nutritional State Modulates the Neural Processing of Visual Motion. Current Biology 24, 890–895 (2014).

13. S. J. Huston, H. G. Krapp, Nonlinear Integration of Visual and Haltere Inputs in Fly Neck Motor Neurons. J. Neurosci. 29, 13097–13105 (2009).

14. J. A. Bender, M. H. Dickinson, A comparison of visual and haltere-mediated feedback in the control of body saccades in Drosophila melanogaster. Journal of Experimental Biology 209, 4597–4606 (2006).

15. A. Dahake, A. L. Stöckl, J. J. Foster, S. P. Sane, A. Kelber, The roles of vision and antennal mechanoreception in hawkmoth flight control. eLife 7, e37606 (2018).

16. D. Rien, R. Kern, R. Kutz, Octopaminergic modulation of a fly visual motion-sensitive neuron during stimulation with naturalistic optic flow. Frontiers in behavioral neuroscience 7 (2013).

17. S. P. Sane, M. J. McHenry, The biomechanics of sensory organs. Integrative and Comparative Biology 49, i8–i23 (2009).

18. J. W. S. Pringle, J. Gray, The gyroscopic mechanism of the halteres of Diptera. Philosophical Transactions of the Royal Society of London. Series B, Biological Sciences 233, 347–384 (1997).

19. M. H. Dickinson, Comparison of Encoding Properties of Campaniform Sensilla on the Fly Wing. Journal of Experimental Biology 151, 245–261 (1990).

20. M. H. Dickinson, Directional Sensitivity and Mechanical Coupling Dynamics of Campaniform Sensilla During Chord-Wise Deformations of the Fly Wing. Journal of Experimental Biology 169, 221–233 (1992).

21. B. H. Dickerson, J. L. Fox, S. Sponberg, Functional diversity from generic encoding in insect campaniform sensilla. Current Opinion in Physiology 19, 194–203 (2021).

22. J. P. Bacon, R. K. Murphey, Receptive fields of cricket giant interneurones are related to their dendritic structure. J Physiol 352, 601–623 (1984).

23. G. A. Jacobs, J. P. Miller, Z. Aldworth, Computational mechanisms of mechanosensory processing in the cricket. Journal of Experimental Biology 211, 1819–1828 (2008).

24. F. G. Barth, Mechanics to pre-process information for the fine tuning of mechanoreceptors. J Comp Physiol A Neuroethol Sens Neural Behav Physiol 205, 661–686 (2019).

25. M. J. Uhrhan, R. J. Bomphrey, H. Lin, Flow sensing on dragonfly wings. Annals of the New York Academy of Sciences 1536, 107–121 (2024).

26. J. C. Tuthill, R. I. Wilson, Mechanosensation and adaptive motor control in insects. Curr Biol 26, R1022–R1038 (2016).

27. T. L. Mohren, T. L. Daniel, S. L. Brunton, B. W. Brunton, Neural-inspired sensors enable sparse, efficient classification of spatiotemporal data. Proc. Natl. Acad. Sci. U.S.A. 115, 10564–10569 (2018).

28. A. I. Weber, T. L. Daniel, B. W. Brunton, Wing structure and neural encoding jointly determine sensing strategies in insect flight. PLoS Comput Biol 17, e1009195 (2021).

29. A. I. Weber, et al., Nonuniform structural properties of wings confer sensing advantages. Journal of The Royal Society Interface 20, 20220765 (2023).

30. F. Reich, D. Warland, R. de Ruyter von Steveninck, W. Bialek, Spikes: Exploring the Neural Code (1997).

31. A. Borst, F. E. Theunissen, Information theory and neural coding. Nat Neurosci 2, 947–957 (1999).

32. G. Rüppell, Kinematic Analysis of Symmetrical Flight Manoeuvres of Odonata. Journal of Experimental Biology 144, 13–42 (1989).

33. J. M. Wakeling, C. P. Ellington, Dragonfly Flight : II. Velocities, Accelerations and Kinematics of Flapping Flight. Journal of Experimental Biology 200, 557–582 (1997).

34. T. L. Hedrick, T. L. Daniel, Flight control in the hawkmoth Manduca sexta: the inverse problem of hovering. J Exp Biol 209, 3114–3130 (2006).

35. J. M. Melis, I. Siwanowicz, M. H. Dickinson, Machine learning reveals the control mechanics of an insect wing hinge. Nature 628, 795–803 (2024).

36. S. M. Walker, et al., In Vivo Time-Resolved Microtomography Reveals the Mechanics of the Blowfly Flight Motor. PLOS Biology 12, e1001823 (2014).

37. S. M. Walker, A. L. R. Thomas, G. K. Taylor, Deformable wing kinematics in the desert locust: how and why do camber, twist and topography vary through the stroke? J R Soc Interface 6, 735–747 (2009).

38. Q. Li, M. Zheng, T. Pan, G. Su, Experimental and Numerical Investigation on Dragonfly Wing and Body Motion during Voluntary Take-off. Sci Rep 8, 1011 (2018).

39. P. J. Simmons, The neuronal control of dragonfly flight. (1977). 10.25911/5d7783acf1631.

40. J. M. McCarthy, S. Watkins, A. Deivasigamani, S. J. John, Fluttering energy harvesters in the wind: A review. Journal of Sound and Vibration 361, 355–377 (2016).

41. C. Koehler, Z. Liang, Z. Gaston, H. Wan, H. Dong, 3D reconstruction and analysis of wing deformation in free-flying dragonflies. Journal of Experimental Biology 215, 3018–3027 (2012).

42. T. L. Hedrick, S. A. Combes, L. A. Miller, Recent developments in the study of insect flight. Can. J. Zool. 93, 925–943 (2015).

43. A. M. Wilson, M. P. McGuigan, A. Su, A. J. van Den Bogert, Horses damp the spring in their step. Nature 414, 895–899 (2001).

44. K. Ishizaka, J. L. Flanagan, Synthesis of voiced sounds from a two-mass model of the vocal cords. The Bell System Technical Journal 51, 1233–1268 (1972).

45. A. J. Hudspeth, How the ear’s works work. Nature 341, 397–404 (1989).

46. W. Cade, Acoustically Orienting Parasitoids: Fly Phonotaxis to Cricket Song. Science 190, 1312–1313 (1975).

47. D. Ko, H.-T. Lin, NBits-Dragonsort. (2024). Deposited 26 May 2024.

48. J. L. Fox, A. L. Fairhall, T. L. Daniel, Encoding Properties of Haltere Neurons Enable Motion Feature Detection in a Biological Gyroscope. Proceedings of the National Academy of Sciences of the United States of America 107, 3840–3845 (2010).

49. A. M. Yarger, J. L. Fox, Single mechanosensory neurons encode lateral displacements using precise spike timing and thresholds. Proc Biol Sci 285, 20181759 (2018).

50. N. I. Fisher, Statistical Analysis of Circular Data (Cambridge University Press, 1993).

51. H. Rajabi, V. Schroeter, S. Eshghi, S. N. Gorb, The probability of wing damage in the dragonfly Sympetrum vulgatum (Anisoptera: Libellulidae): a field study. Biol Open 6, 1290– 1293 (2017).

52. H. Krapp, G. Taylor, J. Humbert, “The mode-sensing hypothesis: Matching sensors, actuators and flight dynamics” in Frontiers in Sensing: From Biology to Engineering, (2012), pp. 101–114.

53. E. Gettrup, Sensory Regulation of Wing Twisting in Locusts. Journal of Experimental Biology 44, 1–16 (1966).

54. B. P. Campbell, J. A. Supple, S. T. Fabian, H.-T. Lin, H. G. Krapp, Adaptive feedforward speed control in Drosophila. [Preprint] (2025). Available at: https://www.biorxiv.org/content/10.1101/2025.03.03.641162v1 [Accessed 5 July 2025].

55. M. Mischiati, et al., Internal models direct dragonfly interception steering. Nature 517, 333–338 (2015).

56. A. A. Makarova, A. A. Polilov, D. B. Chklovskii, Small brains for big science. Current Opinion in Neurobiology 71, 77–83 (2021).

57. F. Song, Y. Yan, J. Sun, Review of insect-inspired wing micro air vehicle. Arthropod Structure & Development 72, 101225 (2023).

58. J. B. Bergmann, D. Moatsou, U. Steiner, B. D. Wilts, Bio-inspired materials to control and minimise insect attachment. Bioinspir. Biomim. 17, 051001 (2022).

59. L. Díaz-García, B. Latham, A. Reid, J. Windmill, Review of the applications of principles of insect hearing to microscale acoustic engineering challenges. Bioinspir. Biomim. 18, 051002 (2023).

60. L. Holden-Dye, R. J. Walker, Invertebrate models of behavioural plasticity and human disease. Brain Neurosci Adv 2, 2398212818818068 (2018).

61. L. Sumathirathne, T. Kim, D. Q. Bower, L. F. Deravi, Cephalopods as a Natural Sensor- Display Feedback System Inspiring Adaptive Technologies. ECS Sens. Plus 2, 023601 (2023).

62. C. Harvey, G. de Croon, G. K. Taylor, R. J. Bomphrey, Lessons from natural flight for aviation: then, now and tomorrow. Journal of Experimental Biology 226, jeb245409 (2023).

63. T. Kim, et al., Wing-strain-based flight control of flapping-wing drones through reinforcement learning. Nat Mach Intell 6, 992–1005 (2024).

64. K. Kubota, H. Tanaka, Machine Learning-Based Wind Classification by Wing Deformation in Biomimetic Flapping Robots: Biomimetic Flexible Structures Improve Wind Sensing. Advanced Intelligent Systems 2400473.

65. V. E. Arriola-Rios, et al., Modeling of Deformable Objects for Robotic Manipulation: A Tutorial and Review. Front Robot AI 7, 82 (2020).

66. F. Nadon, A. J. Valencia, P. Payeur, Multi-Modal Sensing and Robotic Manipulation of Non- Rigid Objects: A Survey. Robotics 7, 74 (2018).

67. D. Rus, M. T. Tolley, Design, fabrication and control of soft robots. Nature 521, 467–475 (2015).

68. J. Montagnat, H. Delingette, N. Ayache, A review of deformable surfaces: topology, geometry and deformation. Image and Vision Computing 19, 1023–1040 (2001).

69. M. Gherlone, P. Cerracchio, M. Mattone, Shape sensing methods: Review and experimental comparison on a wing-shaped plate. Progress in Aerospace Sciences 99, 14– 26 (2018).

70. Z. Ma, J. Choi, H. Sohn, Structural displacement sensing techniques for civil infrastructure: A review. Journal of Infrastructure Intelligence and Resilience 2, 100041 (2023).

71. X. Wang, et al., Recent Progress in Electronic Skin. Advanced Science 2, 1500169 (2015).

72. L. Chen, et al., Spike timing–based coding in neuromimetic tactile system enables dynamic object classification. Science 384, 660–665 (2024).

73. S. Walker, G. Taylor, A semi-empirical model of the aerodynamics of manoeuvring insect flight. Journal of the Royal Society, Interface 18, 20210103 (2021).

74. T. L. Hedrick, Software techniques for two- and three-dimensional kinematic measurements of biological and biomimetic systems. Bioinspir. Biomim. 3, 034001 (2008).

75. J. Blaber, B. Adair, A. Antoniou, Ncorr: Open-Source 2D Digital Image Correlation Matlab Software. Exp Mech 55, 1105–1122 (2015).

76. D. Solav, K. M. Moerman, A. M. Jaeger, K. Genovese, H. M. Herr, MultiDIC: An Open-Source Toolbox for Multi-View 3D Digital Image Correlation. IEEE Access 6, 30520–30535 (2018).

77. S. A. Combes, T. L. Daniel, Into thin air: contributions of aerodynamic and inertial-elastic forces to wing bending in the hawkmoth *Manduca sexta*. Journal of Experimental Biology 206, 2999–3006 (2003).

78. E. Baroth, et al., “IVHM (Integrated Vehicle Health Management) techniques for future space vehicles” in 37th Joint Propulsion Conference and Exhibit, (American Institute of Aeronautics and Astronautics).

79. R. Åke Norberg, The pterostigma of insect wings an inertial regulator of wing pitch. J. Comp. Physiol. 81, 9–22 (1972).

80. S. R. Jongerius, D. Lentink, Structural analysis of a dragonfly wing. Experimental Mechanics 50, 1323–1334 (2010).

81. J. Aljadeff, B. J. Lansdell, A. L. Fairhall, D. Kleinfeld, Analysis of Neuronal Spike Trains, Deconstructed. Neuron 91, 221–259 (2016).

82. S. M. Kay, Modern spectral estimation: theory and application (Prentice Hall, 1988).

83. C. E. Shannon, A mathematical theory of communication. The Bell System Technical Journal 27, 379–423 (1948).

84. F. H. Eeckman, Analysis and Modeling of Neural Systems (Springer US, 1992).

85. H. Clague, F. Theunissen, J. P. Miller, Effects of adaptation on neural coding by primary sensory interneurons in the cricket cercal system. J Neurophysiol 77, 207–220 (1997).

86. D. K. Warland, P. Reinagel, M. Meister, Decoding visual information from a population of retinal ganglion cells. J Neurophysiol 78, 2336–2350 (1997).

87. R. Wessel, C. Koch, F. Gabbiani, Coding of time-varying electric field amplitude modulations in a wave-type electric fish. J Neurophysiol 75, 2280–2293 (1996).

88. M. Juusola, A. S. French, The Efficiency of Sensory Information Coding by Mechanoreceptor Neurons. Neuron 18, 959–968 (1997).

89. F. Theunissen, J. C. Roddey, S. Stufflebeam, H. Clague, J. P. Miller, Information theoretic analysis of dynamical encoding by four identified primary sensory interneurons in the cricket cercal system. J Neurophysiol 75, 1345–1364 (1996).

90. G. T. Buraĉas, A. M. Zador, M. R. DeWeese, T. D. Albright, Efficient Discrimination of Temporal Patterns by Motion-Sensitive Neurons in Primate Visual Cortex. Neuron 20, 959– 969 (1998).

91. R. A. DiCaprio, C. P. Billimoria, B. C. Ludwar, Information rate and spike-timing precision of proprioceptive afferents. J Neurophysiol 98, 1706–1717 (2007).

92. W. Bialek, F. Rieke, R. R. De Ruyter Van Steveninck, D. Warland, Reading a Neural Code. Science 252, 1854–1857 (1991).

93. H. Wolf, The Locust Tegula: Significance for Flight Rhythm Generation, Wing Movement Control and Aerodynamic Force Production. Journal of Experimental Biology 182, 229–253 (1993).

94. M. A. Frye, Effects of stretch receptor ablation on the optomotor control of lift in the hawkmoth Manduca sexta. J Exp Biol 204, 3683–3691 (2001).

95. E. Marder, A. L. Taylor, Multiple models to capture the variability in biological neurons and networks. Nat Neurosci 14, 133–138 (2011).

96. J. A. Supple, et al., Binocular Encoding in the Damselfly Pre-motor Target Tracking System. (2020). 10.17863/CAM.48011.

## SI References

1. E. Gettrup, Sensory Regulation of Wing Twisting in Locusts. Journal of Experimental Biology 44, 1–16 (1966).

2. H. Wolf, The Locust Tegula: Significance for Flight Rhythm Generation, Wing Movement Control and Aerodynamic Force Production. Journal of Experimental Biology 182, 229–253 (1993).

3. M. A. Frye, Effects of stretch receptor ablation on the optomotor control of lift in the hawkmoth Manduca sexta. J Exp Biol 204, 3683–3691 (2001).

4. E. Marder, A. L. Taylor, Multiple models to capture the variability in biological neurons and networks. Nat Neurosci 14, 133–138 (2011).

5. J. A. Supple, et al., Binocular Encoding in the Damselfly Pre-motor Target Tracking System. (2020). 10.17863/CAM.48011.

6. K. D. Longden, T. Muzzu, D. J. Cook, S. R. Schultz, H. G. Krapp, Nutritional State Modulates the Neural Processing of Visual Motion. Current Biology 24, 890–895 (2014).

7. S. J. Huston, H. G. Krapp, Nonlinear Integration of Visual and Haltere Inputs in Fly Neck Motor Neurons. J. Neurosci. 29, 13097–13105 (2009).

8. D. Rien, R. Kern, R. Kutz, Octopaminergic modulation of a fly visual motion-sensitive neuron during stimulation with naturalistic optic flow. Frontiers in behavioral neuroscience 7 (2013).

